# Emergence of robust nucleosome patterns from an interplay of positioning mechanisms

**DOI:** 10.1101/431445

**Authors:** Johannes Nuebler, Michael Wolff, Benedikt Obermayer, Wolfram Möbius, Ulrich Gerland

## Abstract

Proper positioning of nucleosomes in eukaryotic cells is determined by a complex interplay of factors, including nucleosome-nucleosome interactions, DNA sequence, and active chromatin remodeling. Yet, characteristic features of nucleosome positioning, such as gene-averaged nucleosome patterns, are surprisingly robust across perturbations, conditions, and species. Here, we explore how this robustness arises despite the underlying complexity. We leverage mathematical models to show that a large class of positioning mechanisms merely affects the quantitative characteristics of qualitatively robust positioning patterns. We demonstrate how statistical positioning emerges as an effective description from the complex interplay of different positioning mechanisms, which ultimately only renormalize the model parameter quantifying the effective softness of nucleosomes. This renormalization can be species-specific, rationalizing a puzzling discrepancy between the effective nucleosome softness of *S. pombe* and *S. cerevisiae.* More generally, we establish a quantitative framework for dissecting the interplay of different nucleosome positioning determinants.

## INTRODUCTION

Eukaryotic DNA is condensed into chromatin in a hierarchy of spatial organization and compaction levels. This organization is dynamic on all scales, from the large-scale organization of the genome (Bonev and Cavalli, 2016; Fraser et al., 2015) down to the positions of individual nucleosomes, the smallest subunit of chromatin, which consists of 147 base pairs of DNA wrapped around an octamer of histone proteins (Luger et al., 1997). Since nucleosomes restrict accessibility of other factors to DNA, their positioning is essential for gene regulation (Bai and Morozov, 2010; Korber and Barbaric, 2014; Lam et al., 2008; Teif et al., 2013; Venkatesh et al., 2013).

Genome wide mapping of nucleosome positions (Voong et al., 2017) reveals gene-specific nucleosome patterns, which survive the average over many cells inherent to these methods. Known determinants of nucleosome positions (Hughes and Rando, 2014; Struhl and Segal, 2013) include the DNA sequence (Kaplan et al., 2009; Segal et al., 2006), competition with other DNA binding proteins (Ozonov and van Nimwegen, 2013), ATP-dependent chromatin remodeling enzymes that can relocate, modify or evict nucleosomes (Bartholomew, 2014; Clapier et al., 2017; Mueller-Planitz et al., 2013; Zhou et al., 2016), RNA polymerase and the DNA replication machinery (Radman-Livaja et al., 2011; Weiner et al., 2010), and interactions between the nucleosomes themselves, which can partially invade each other, due to unwrapping of DNA from the histone core (Chereji and Morozov, 2014; Engeholm et al., 2009). However, a complete understanding of how gene specific nucleosome patterns emerge from this interplay remains elusive.

The complex gene-specific nucleosome patterns yield a simple, characteristic pattern when averaged over many genes (Yuan et al., 2005), with a depleted promoter region followed by a downstream array. The nucleosomes within this array display a degree of variability in their positions that increases with distance from the promoter, as reflected in the oscillatory nucleosome density with peaks of decaying amplitude and increasing widths, cf. Fig. 1. The qualitative shape of this consensus pattern is universal across species with the nucleosome peak to peak distance varying from 150 base pairs (bp) in the yeast *S. pombe* (Lantermann et al., 2010; Moyle-Heyrman et al., 2013) to more than 200 bp in humans (Schones et al., 2008; Valouev et al., 2011). Furthermore, the shape is surprisingly robust against a multitude of perturbations: introduction of foreign DNA into the genome (Hughes et al., 2012), substitution of remodeler-encoding genes by non-endogenous variants and removal of H1 linker histones (Hughes and Rando, 2016), reduction of the overall histone abundance (Celona et al., 2011; Gossett and Lieb, 2012), different growth conditions (Kaplan et al., 2009), and diamide stress (Weiner et al., 2015) all affect the oscillatory consensus pattern only mildly. The passage of RNA polymerase also appears to have only a minor influence on the gene-averaged nucleosome pattern (Bintu et al., 2011; Radman-Livaja et al., 2011; Weiner et al., 2010), given that the pattern is only weakly dependent on transcription rate in yeast (Chereji and Morozov, 2015; Weiner et al., 2010). However, some amount of transcription may be required to establish the pattern, since silent genes show very weak oscillations in *Drosophila melanogaster* (Chereji et al., 2016). Substantial changes in the gene-averaged nucleosome pattern have been observed shortly after replication (Fennessy and Owen-Hughes, 2016; Vasseur et al., 2016) and in some remodeler deletion strains, which display reduced positioning oscillations (Gkikopoulos et al., 2011; Ocampo et al., 2016).

**FIG. 1:**
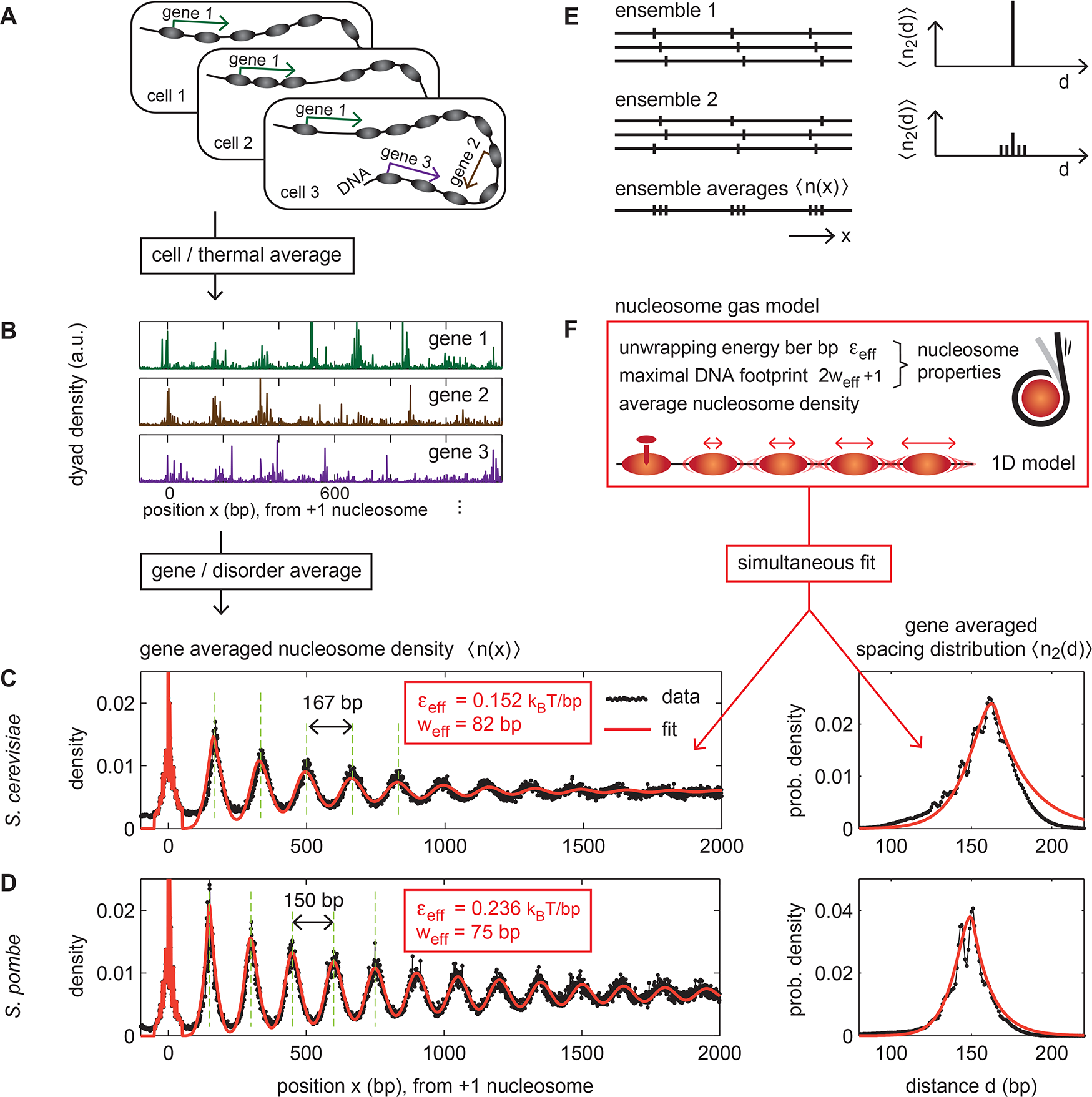
Surprising compatibility between nucleosome positioning data and soft-core nucleosome gas model. Characteristic patterns emerge when nucleosome positions are averaged across cells (A) and genes (B). (C,D) Black dots show the gene averaged nucleosome density (left) and the spacing distribution (distribution of dyad-to-dyad distances, right) in *S. cerevisiae* and *S. pombe* from chemical cleavage data (Brogaard et al., 2012; Moyle-Heyrman et al., 2013). The average is taken over genes longer than 2500 bp, aligned by the most likely position of their first nucleosome. (E) The spacing distribution provides additional statistical information about the nucleosome configurations of individual cells that is not contained in the nucleosome density: The two ensembles have the same cell-averaged density profile 〈*n*(*x*)〉, but different spacing distributions 〈*n*_2_(*d*)〉. (F) A physical model for nucleosome positioning: the nucleosome gas model with a fixed barrier. Two parameters characterize the nucleosome properties: *w*_eff_ parametrizes the maximal DNA footprint, and *ε*_eff_ is the effective nucleosome stiffness, i.e. the effective energy cost for unwrapping nucleosomal DNA. This model accurately describes the gene averaged nucleosome density 〈*n*(*x*)〉 and spacing distribution 〈*n*_2_(*d*)〉 simultaneously: the red curves in (C,D) show the best fit to the data. The estimated nucleosome stiffness and footprint differ substantially between *S. cerevisiae* and *S. pombe.* We show here that these differences can emerge from species-specific positioning mechanisms rather than pointing to different properties of the histones, which are highly conserved across species.

Why are gene-averaged nucleosome patterns so universal (across species) and robust (against perturbations)? An important step towards addressing this fundamental question was made by Kornberg and Stryer (Kornberg and Stryer, 1988), who suggested that nucleosome patterns could be described by a simple model, known as the ‘Tonks gas’ in physics (Tonks, 1936). The model assumes that nucleosome arrangements along the DNA correspond to an equilibrium ensemble of extended, non-overlapping particles positioned at a certain average density along a one-dimensional substrate. A perturbation in this gas, created for instance by a repulsive barrier, then induces oscillatory patterns in the average positions of the adjacent particles, which seemed consistent with the ladders observed in gels after nuclease digestion (Kornberg and Stryer, 1988). Later, when whole-genome nucleosome mapping became feasible, the model could be compared to the average nucleosome patterns in the vicinity of promoters (Mavrich et al., 2008). It was shown that the patterns downstream of yeast promoters are in good agreement with the model, if the perturbation is assumed to be created by a mechanism that holds the first nucleosome in a fixed position, whereas the patterns upstream of the promoters are compatible with a repulsive barrier (Möbius and Gerland, 2010).

While this “nucleosome gas” model originally treated nucleosomes as hard particles with footprints that cannot overlap on DNA, subsequent analyses took into account the “softness” of nucleosomes arising from transient DNA unwrapping from the histone core (Chereji and Morozov, 2014; Möbius et al., 2013). With this addition, the nucleosome gas model was able to provide a unified effective description of the gene-averaged nucleosome patterns across different *Ascomycota* fungi (Möbius et al., 2013). However, while this model can describe the data, it does not suffice to explain it. *A priori,* the effectiveness of such a simple description even seems unreasonable, since it neither accounts for the sequence specificity of the histone-DNA interaction, nor the action of remodeling enzymes. The resolution of this conundrum is our primary goal here.

We first illustrate the surprising effectiveness of the simple nucleosome gas model by showing that it simultaneously describes two different properties of the nucleosome patterns, the gene averaged nucleosome density and the spacing distribution, in both *S. cerevisiae* and *S. pombe.* This provides the basis for our subsequent analysis of how the interplay of various nucleosome positioning mechanisms can produce the robust nucleosome gas features. Towards this end, we design a class of nucleosome positioning models that explicitly incorporate the effects of active remodeling and DNA sequence-dependent nucleosome stability. We leverage the modeling framework to demonstrate that many (but not all) of the considered mechanisms are compatible with the simple nucleosome gas model, in the sense that they merely modify (or “renormalize”) the apparent softness of the nucleosomes. As a consequence, the resulting gene-averaged patterns display a high degree of universality, because their qualitative shape is not produced by a fine-tuned balance of specific mechanisms, but emerges robustly from the interplay of different positioning mechanisms. We suggest that a particular renormalization effect in *S. pombe* can explain why nucleosomes have a different apparent softness in this species compared to *S. cerevisiae* and related species, even though the underlying histone proteins are highly conserved. Finally, we use our models to infer which nucleosome positioning scenarios are incompatible with the available data, and discuss which novel types of experimental data would provide more effective discrimination between different scenarios.

## RESULTS

To study the interplay of nucleosome positioning mechanisms, one would ideally monitor the arrangements of groups of nucleosomes in individual cells over time. Such dynamical information would directly reveal the effects of positioning mechanisms, and different mechanisms could be disentangled with appropriate mutants. Instead, the established whole-genome nucleosome mapping techniques provide statistical information, due to the ensemble average over large numbers of cells that is inherent to these techniques (Fig. 1A). In the case of MNase-seq data, a sequencing read reflects the DNA footprint of a single nucleosome in one of these cells. In contrast, the chemical cleavage technique (Brogaard et al., 2012) yields sequencing reads of the DNA connecting the midpoints (dyads) of two nucleosomes. Both techniques permit the extraction of nucleosome density profiles, *n*(*x*), i.e. histograms of nucleosome dyad positions along the DNA (Fig. 1B). Gene-averaged nucleosome density profiles, 〈*n*(*x*)〉, yield the consensus nucleosome pattern for a set of genes, e.g. see Fig. 1C,D (left panels) for the *S. cerevisiae* and *S. pombe* patterns, respectively (here, *x* denotes the distance in basepairs to the +1 nucleosome, see caption and ‘Methods’). Such consensus patterns haven proven to be very reproducible, and are commonly used as quantitative signatures of the interplay between different nucleosome positioning mechanisms in different species or mutants (Gkikopoulos et al., 2011; Hughes and Rando, 2016; Tsankov et al., 2010; Zhang et al., 2011).

The chemical cleavage technique (Brogaard et al., 2012) also permits to extract another type of statistical information, the nucleosome spacing distribution 〈*n*_2_(*d*)〉, i.e. the distribution of distances *d* between the dyads of neighboring nucleosomes, shown in Fig. 1C,D (right panels). Importantly, the spacing distribution provides additional statistical information about the nucleosome configurations of individual cells, which is not contained in the nucleosome density pattern. This is illustrated in Fig. 1E, by showing two ensembles of nucleosome configurations that yield the same ensemble-averaged nucleosome density profiles 〈*n*(*x*)〉, but different spacing distributions 〈*n*_2_(*d*)〉. We therefore treat both of these statistical characteristics as quantitative signatures of the interplay between different nucleosome positioning mechanisms.

### Robust statistical characteristics of nucleosome positioning

To illustrate the conundrum that motivated this study, we analyze the quantitative signatures 〈*n*(*x*)〉 and 〈*n*_2_(*d*)〉 in *S. cerevisiae* and *S. pombe,* based on existing chemical cleavage maps (Brogaard et al., 2012; Moyle-Heyrman et al., 2013). As seen in Fig. 1C,D (black dots), the data displays qualitatively similar signatures in these evolutionarily distant yeast species. The gene-averaged nucleosome density profiles show pronounced oscillations, with amplitudes that decay with increasing distance from the first nucleosome (the “+1” nucleosome) of the genes, while the spacing distributions display a cusp-like peak with a fine-grained substructure. However, on a quantitative level, the signatures differ significantly between the two species. The peak-to-peak distance in 〈*n*(*x*)〉 is much smaller in *S. pombe* (150 bp) than in *S. cerevisiae* (167 bp), and the oscillations persist over a longer range in *S. pombe*. Furthermore, the spacing distribution is considerably wider in *S. cerevisiae* than in *S. pombe*, and peaked at a larger average spacing.

*S. cerevisiae* is the best-studied representative from a group of *Ascomycota* fungi, for which the gene-averaged nucleosome density profiles 〈*n*(*x*)〉 have previously been analyzed with the “nucleosome gas” model (Möbius et al., 2013). This analysis revealed that the precise shape of 〈*n*(*x*)〉, as well as its species-to-species variation within this group, can be described within the same model. Within this model, the differences between the profiles of different species arise purely as a consequence of the different nucleosome packing densities, or, equivalently, different average linker lengths, as opposed to different nucleosome properties. A conceptually very similar model was also used to interpret the spacing distribution of *S. cerevisiae,* again finding surprisingly good agreement(Chereji and Morozov, 2014). The study showed that the model can even capture the fine-grained structure of 〈*n*_2_(*d*)〉, by taking into account that nucleosomal DNA tends to unwrap from the nucleosome core in discrete steps of one helical turn.

Here, we adopt the “nucleosome gas” model (Möbius et al., 2013), which describes the nucleosome properties by two effective biophysical parameters, see Fig. 1F. The softness of nucleosomes, i.e. their ability to invade each others DNA footprint (Engeholm et al., 2009), is taken into account in a coarse grained way by assuming an energetic cost *ε*_eff_ per base pair to unwrap DNA from the histone core. Furthermore, the reach of the repulsive interaction between nucleosomes is parameterized by the effective nucleosome footprint radius *w*_eff_, corresponding to a maximal footprint of 2*w*_eff_ + 1 base pairs, which is expected to exceed the canonical 147 base pair length of nucleosomal DNA, due to steric constraints arising when two nucleosomes are very close. The resulting effective nucleosome-nucleosome interaction potential is derived by summing the statistical weights of all unwrapping states compatible with a given nucleosome separation *d* (measured dyad-to-dyad), see Supplement for a detailed description of all Methods.

In contrast to prior studies, we test whether the model is able to simultaneously describe both signatures, 〈*n*(*x*)〉 and 〈*n*_2_(*d*)〉, with the same parameter values. The red lines in Fig. 1C,D show the resulting best fit. For *S. cerevisiae,* the fit yields an effective interaction radius of *w*_eff_ = 82 bp and an effective unwrapping cost of *ε*_eff_ = 0.152 *k_B_T*/bp. This is consistent with the previous results for a group of *Ascomycota* fungi that included *S. cerevisiae,* where *w*_eff_ = 83 bp, *ε*_eff_ = 0.153 *k_B_T*/bp were obtained from the density patterns alone (Möbius et al., 2013). Here we find that the spacing distribution is compatible with the same nucleosome gas model and parameters. The visible discrepancy in 〈*n*_2_(*d*)〉 for large spacings is expected, since long sequencing reads are suppressed in the data, see Supplement.

For *S. pombe,* the simultaneous fit provides an even better description of the density profiles and the spacing distribution, see Fig. 1D. However, the corresponding parameter values, *w*_eff_ = 75 bp and *ε*_eff_ = 0.236 *k_B_T*/bp, deviate markedly from those for *S. cerevisiae* and the previously studied group of *Ascomycota* fungi. Fig. S1A-C shows that the fitting parameters are well constrained for both species, aided by the inclusion of the spacing distribution in the fit. As an additional test of the consistency of the model, we also reverse engineered the nucleosome-nucleosome interaction directly from the experimentally measured spacing distributions, finding good agreement with our best fit model, see Supplement and Fig. S1D,E.

Taken together, these results support the nucleosome gas model as an effective description of gene averaged *in vivo* nucleosome data across evolutionarily distant yeast species. This implies that the statistical characteristics of the nucleosome gas are robust features of nucleosome positioning. How does this robustness arise despite the underlying complexity of nucleosome positioning mechanisms? In the remainder of this article, we seek to resolve this conundrum. A second question triggered by the above results is why the effective nucleosome stiffness *ε*_eff_ is so different between *S. pombe* and *S. cerevisiae.* We will see that these questions are indeed related: the nucleosome properties observed in the gene average are not the bare biophysical nucleosome properties, because they are renormalized by additional positioning determinants – and this renormalization can be different in different species.

To address these questions, we probe the effect of several simplified, but biologically motivated nucleosome positioning mechanisms in the context of the nucleosome gas model. We use the general approach outlined in Fig. 2. First, we compute the gene-averaged nucleosome density 〈*n*(*x*)〉 and the internucleosomal spacing distribution 〈*n*_2_(*d*)〉 for a modified nucleosome gas model that features an additional positioning determinant, e.g. DNA sequence specificity or a remodeling mechanism. Then, we fit 〈*n*(*x*)〉 and 〈*n*_2_(*d*)〉 by *only* the nucleosome gas model. Thereby we determine two things. First, the goodness of the fit, which measures how compatible the specific positioning determinant is with an effective description by the nucleosome gas model. Second, we determine to what extent the effective parameters *ε*_eff_ and *w*_eff_ differ from the bare parameters *ε* and w. In the following, we apply this framework to several different positioning mechanisms.

**FIG. 2:**
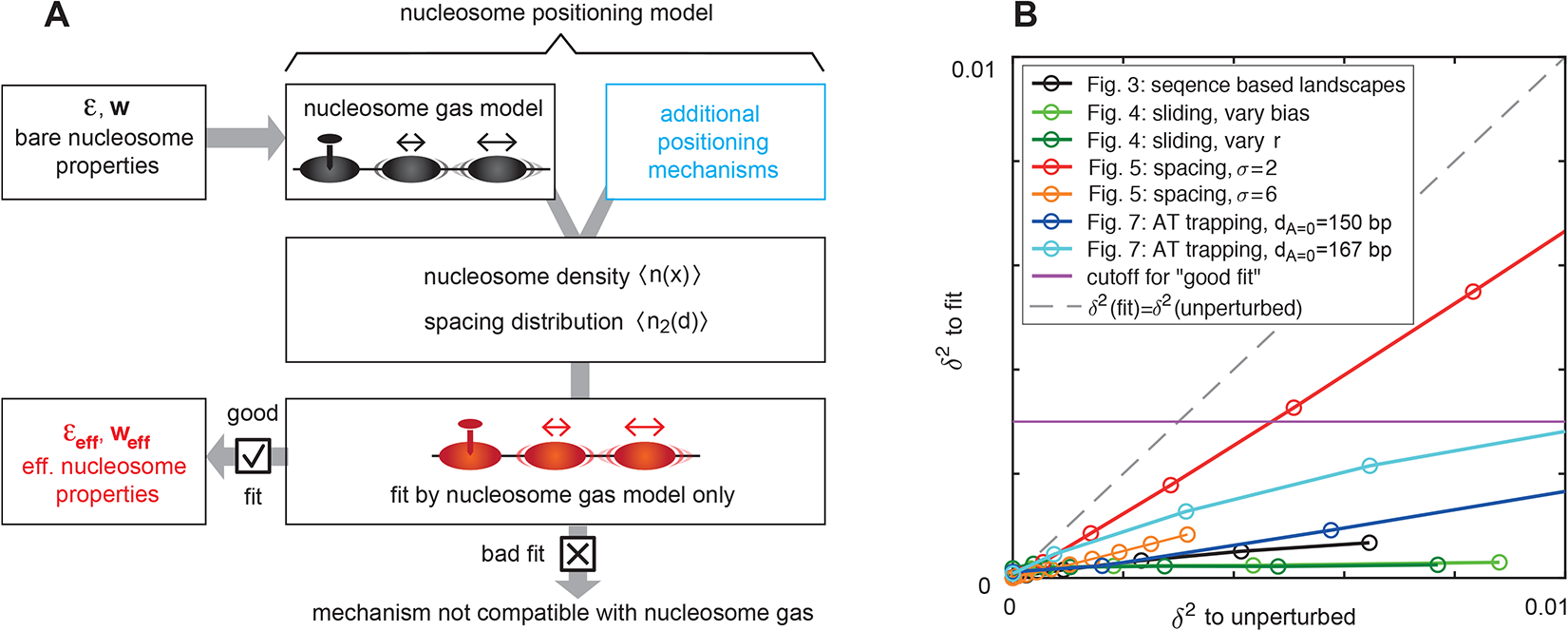
General framework to analyze the effect of specific nucleosome positioning mechanisms on the nucleosome gas. (A) We compute the nucleosome density 〈*n*(*x*)〉 and spacing distribution 〈*n*_2_(*d*)〉 resulting from the interplay of the “nucleosome gas model” with bare nucleosome parameters *ε* and *w* (stiffness and size) and additional positioning mechanisms. Then, we determine if 〈*n*(*x*)〉 and 〈*n*_2_(*d*)〉 can be described by the nucleosome gas model alone. If that is the case (fit good), the effective nucleosome properties *ε*_eff_ and *w*_eff_ may differ from the bare ones. (B) To assess if a nucleosome positioning model can be effectively described by the nucleosome gas model we compare the fit error, δ^2^ (to fit) to the perturbation by the additional positioning mechanism, δ^2^(to unperturbed), namely the squared deviation of 〈*n*(*x*)〉 and 〈*n*_2_(*d*)〉 in the presence of the positioning mechanism from the same quantities without it. For most considered positioning mechanisms, to be discussed below, the fit error is much smaller than the perturbation, indicating good compatibility with the nucleosome gas model. Only for the spacing mechanism with accurate spacing (*σ* = 2 bp) we obtain a fit error that of similar magnitude as the perturbation, indicating that such accurate nucleosome spacing is not compatible with the nucleosome gas model.

### DNA sequence specific positioning leads to effective nucleosome softening

We first ask how gene-specific nucleosome positioning encoded in the DNA sequence affects gene-averaged patterns. We use energy landscapes to account for DNA sequence specificity (Fig. 3B,C). Various models for such energy landscapes have been developed (Tolkunov and Morozov, 2010), e.g. based on DNA elasticity (Eslami-Mossallam et al., 2016; Morozov et al., 2009; Tolstorukov et al., 2007), machine learning (Kaplan et al., 2009), or simple rules (van der Heijden et al., 2012). Here, our goal is not to assess the predictive power of these models, but rather to describe their generic effects on gene-averaged data. We will find that in this context the main parameter is the “positioning power” of energy landscapes, which scales with their standard deviation *σ* (Fig. 3C). Specifically, we use a DNA elasticity based model (Morozov et al., 2009). Exemplary landscapes u(x) for *S. cerevisiae* genes are shown in Fig. 3C (cyan). We consider only genes longer than 2500 bp to avoid interference with positioning effects at gene ends.

**FIG. 3:**
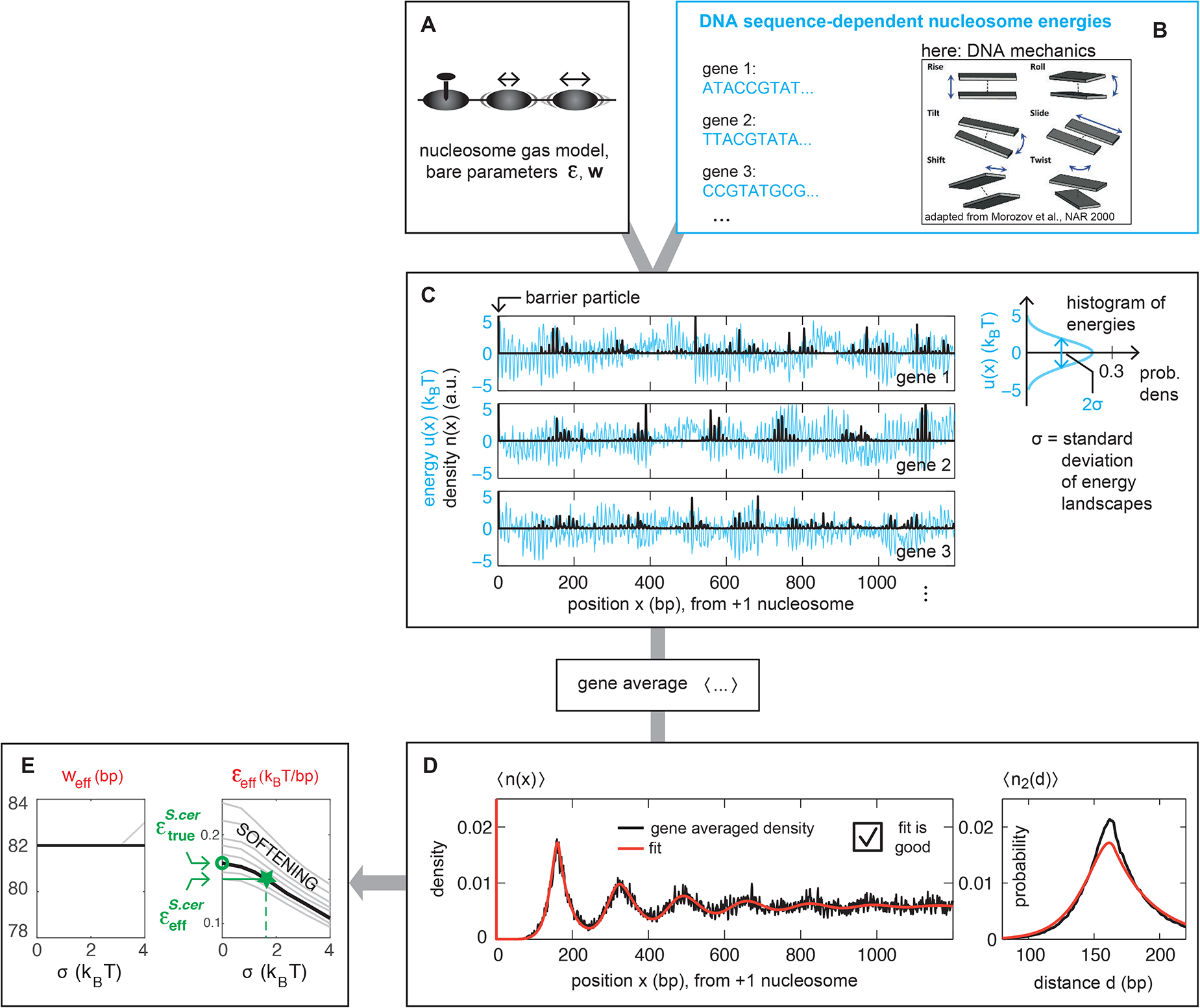
Effective nucleosome softening by DNA sequence-dependent nucleosome positioning. Following the general framework of Fig. 2, we use (A) the nucleosome gas model with bare nucleosome parameters *ε* (stiffness) and *w* (size), together with (B) DNA sequence specific nucleosome positioning energy landscapes to compute the nucleosome density *n*(*x*) and internucleosomal spacing distribution *n*_2_(*d*) on each gene (C, only *n*(*x*) is shown). The gene averaged quantities (D) can be reproduced by the nucleosome gas model *without* landscapes. The fit parameters *ε*_eff_ and *w*_eff_ are the effective nucleosome properties, accounting for the effect of landscapes in the gene average. (E) This procedure is repeated for different standard deviations *σ* of the landscapes (grey lines) and different bare nucleosome stiffnesses *ε*. The flow line intersecting the experimentally observed 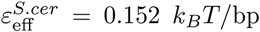 at the best estimate for *σ* = 1.56 belongs to the bare parameter 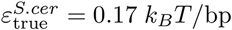 (thick black line). The nucleosome footprint parameter *w* is mostly unaffected by energy landscapes.

We compute the nucleosome density and spacing distribution on each of the obtained landscapes as follows: we fix a barrier particle at the consensus position of the first nucleosome and compute the downstream densities and spacing distributions exactly using the transfer matrix method for given values of the nucleosomes properties *w* and *ε* (Ssupplement). The resulting nucleosome densities (Fig. 3C, black) are strongly influenced by the energy landscapes and thus differ between genes. From these, we compute 〈*n*(*x*)〉 and 〈*n*_2_(*d*)〉, where 〈〉 is the average across genes (Fig. 3D, black). We point out that it is crucial to *first* compute the nucleosome density on each gene and *then* take the average, not *vice versa:*

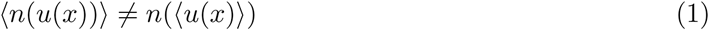

Indeed, the gene-averaged landscape 〈*u*(*x*)〉 is almost flat (*σ* = 0.056 *k_B_T* for our set of 914 long genes in *S. cerevisiae,* Suppl. Fig. S3E), which indicates that there are no sequence encoded positioning clues in coding regions on average according to the used landscape model.

Next, we determine to what extent the nucleosome gas model is compatible with the gene averaged density and spacing distribution by fitting the model to 〈*n*(*x*)〉 and 〈*n*_2_(*d*)〉 simultaneously (Fig. 3D, red, see Suppl. Fig. S2 for details). We find that the gene averaged quantities can be surprisingly well reproduced by this simple model with renormalized nucleosome properties *ε*_eff_ and *w*_eff_. We also find that nucleosomes in the gene average appear softer than the bare particles, *ε*_eff_ < *ε*, while the maximal footprint size is almost unaffected, *w*_eff_ ≈ *w*. The degree of effective nucleosome softening depends on the average landscape amplitude *σ* (Fig. 3E). We determine a realistic value of *σ* from the peakedness of the nucleosome density on single *S. cerevisiae* genes to be *σ^S^*·*^cer^* = 1.56 *k_B_T* (green dashed line in Fig. 3E, see also Suppl. Fig. S3A).

Finally, we determine how stiff individual nucleosomes are “truly”, given that they appear softened by energy landscapes. To do so, we start from the nucleosome stiffness which fits the *S. cerevisiae* data, and from the above estimate for the landscape amplitude, i.e. from the point 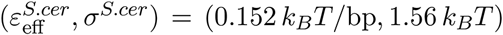 (Fig. 3E, green star). We then trace back the renormalization flow and find the bare nucleosome stiffness 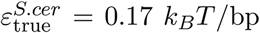 (green circle). Our results thus suggest that this is the true average energetic cost of unwrapping one bp of DNA from a nucleosome in *S. cerevisiae*.

In order to show that the observed robustness and parameter renormalization are not a peculiarity of the chosen energy landscape model we repeated the above steps with uncorrelated landscapes, where for every lattice site an energy is drawn at random from a Gaussian distribution with standard deviation *σ*. The renormalization flow is very similar (Suppl. Fig. S3D) which indicates that the effective nucleosome softening is a generic phenomenon of averaging over nucleosome positioning landscapes.

We have found that positioning by gene specific energy landscapes is “renormalizable”, i.e. the gene averaged quantities can be reproduced without landscapes by altered nucleosome properties. We point out that it is easy to come up with landscapes that do not belong to this renormalizable class. For example, by specific variations in the landscape amplitude *σ* or by introducing specific correlations we obtain gene averaged patterns that are not reproduced by the nucleosome gas model (Fig. S4). We thus conclude that *S. cerevisiae* energy landscapes are random enough to be renormalizable.

### Remodeler effects: directional sliding

Like DNA sequence, nucleosome remodeling enzymes (“remodelers”) are a crucial determinant of nucleosome positioning. They are required for proper nucleosome spacing (Krietenstein et al., 2016; Lieleg et al., 2015; Zhang et al., 2011), and are known to relocate nucleosomes along the DNA using energy from ATP hydrolysis (Flaus and Owen-Hughes, 2011; Mueller-Planitz et al., 2013). It is therefore remarkable that the emerging *in vivo* patterns are well described by the equilibrium nucleosome gas model. Here, we rationalize this agreement by showing that many remodeling mechanisms are indeed “renormalizable”.

We first ask how directional sliding of nucleosomes by remodelers affects positioning. Our minimal model is guided by two observations. First, *in vitro* reconstitution experiments result in stereotypic nucleosome array patterns only in the presence of ATP (Zhang et al., 2011). This indicates that remodelers mobilize nucleosomes that are otherwise kinetically frozen, which is also supported by direct observations of strongly increased DNA unwrapping rates upon remodeler binding (Singh et al., 2018). Second, remodelers can displace nucleosomes in successive steps in the same direction (Blosser et al., 2009). Here, we consider two types of remodelers, upstream (U) and downstream (D) sliders, which preferentially move nucleosomes upstream/downstream along the DNA (they could represent the same protein binding nucleosomes in different orientations). Both bind and unbind nucleosomes with rates 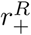 and 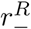, respectively (Fig. 4A). Remodeler bound nucleosomes perform a biased random walk with a rate parameter *r* and a bias parameter *γ*, namely *r_u_,_d_* = *re*^±γ/2^ (see Fig. 4A and Supplement). The parameters *r* and *γ* account for general mobilization and directionality, respectively. We measure the remodeler effectiveness by its processivity *p*, the mean displacement during its dwell time in the absence of interactions with other particles (Supplement). Nucleosome interactions alter the binding and sliding rates, and simulations are performed with a kinetic Monte Carlo scheme (Supplement).

**FIG. 4:**
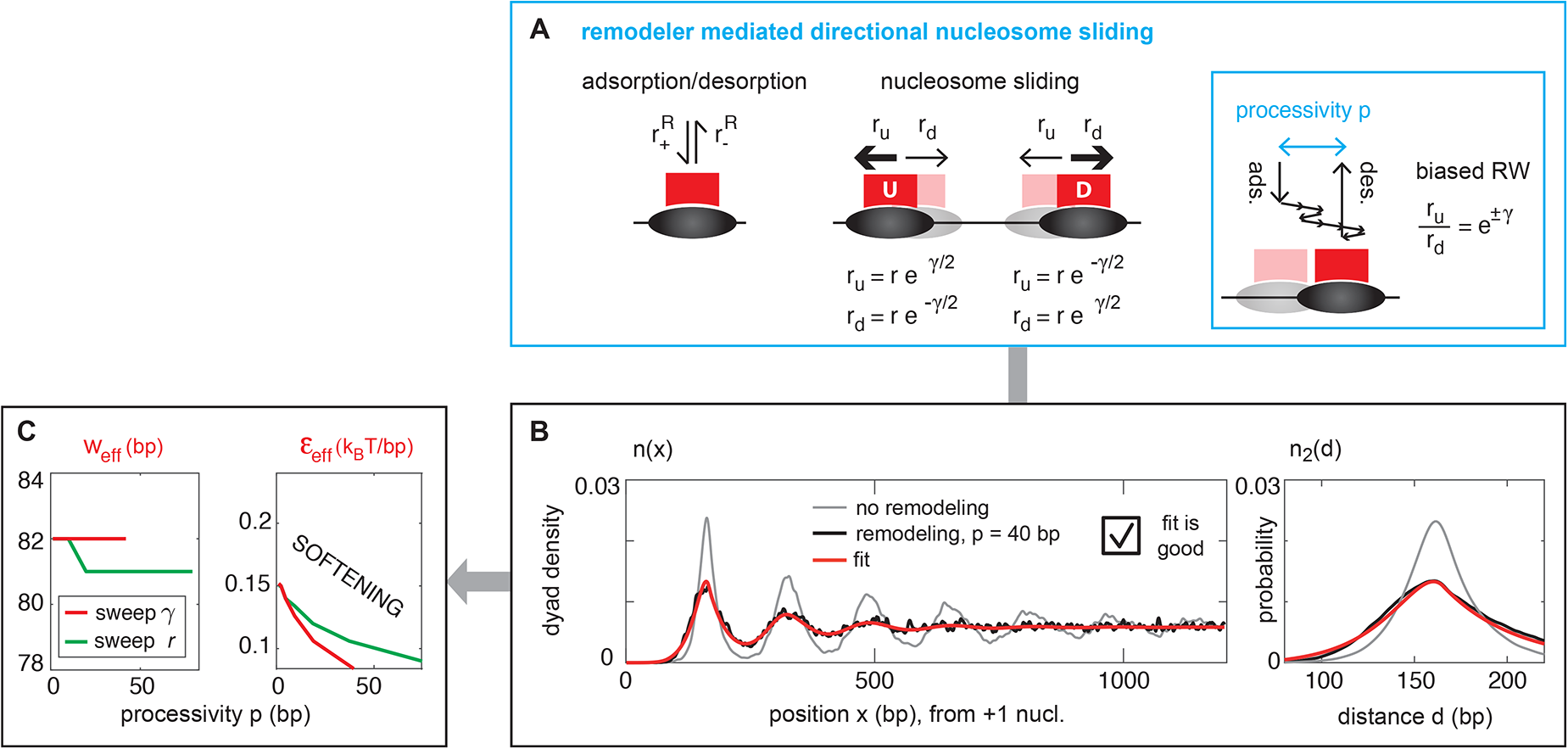
Effective nucleosome softening by remodeler mediated directional sliding. (A) We model nucleosome sliding in the presence of equal amounts of upstream (U) and downstream (D) remodelers, which can bind to nucleosomes and then mediate a random walk that is biased in the upstream/downstream direction. Sliding rates as shown are parameterised by the rate *r* and the bias *γ*. We measure the effectiveness of a remodeler by its processivity, i.e. the average unobstructed displacement during the remodeler residence time (see Supplement). (B) Nucleosome sliding strongly reduces nucleosome density oscillations. Yet, the fit by the nucleosome gas model is excellent. (C) The effective nucleosome properties show a softening for increasing remodeler processivity, which we alter by sweeping the directional bias *γ* or the rate *r*.

We find that nucleosome sliding strongly reduces the characteristic oscillations in the nucleosome density *n*(*x*) and broadens the spacing distribution *n*_2_(*d*) (Fig. 4B, see Fig. S5 for a parameter sweep). Yet, both quantities are excellently compatible with the nucleosome gas model (red vs black lines in Fig. 4B). The effective nucleosome stiffness *ε*_eff_ decreases for increasing processivity *p* (Fig. 4C). This is expected, since our sliding remodelers promote nucleosomes invading each other. The softening is more pronounced when the processivity is altered via the bias parameter *γ* instead of the rate parameter *r* because the bias is more important in overcoming interactions with neighboring nucleosomes (see Supplement). In conclusion, we found that remodeler mediated directional nucleosome sliding is excellently compatible with the nucleosome gas model and leads to effective nucleosome softening.

### Remodeler effects: nucleosome attraction and spacing

Next, we address remodeler mediated nucleosome attraction and spacing. *In vitro* (Lieleg et al., 2015; Zhang et al., 2011) and *in vivo* (Celona et al., 2011; Gossett and Lieb, 2012) experiments have shown that in the presence of remodelers nucleosome spacing does not change even when the nucleosome abundance is strongly reduced. This is incompatible with the nucleosome gas model, which predicts that the average nucleosome spacing increases for reduced density. Thus, remodeler mediated attraction was proposed to maintain a fixed spacing in gene-coding regions (Lieleg et al., 2015; Möbius et al., 2013).

We implement attraction towards a fixed spacing by superimposing the repulsive soft-core nucleosome interaction with an attractive potential well centred around the *S. cerevisiae* spacing of *d*_0_ = 167 bp (Fig. 5A). Its standard deviation *σ* parametrizes the remodeler’s spacing accuracy and the depth *a* the spacing activity (see Supplement).

**FIG. 5:**
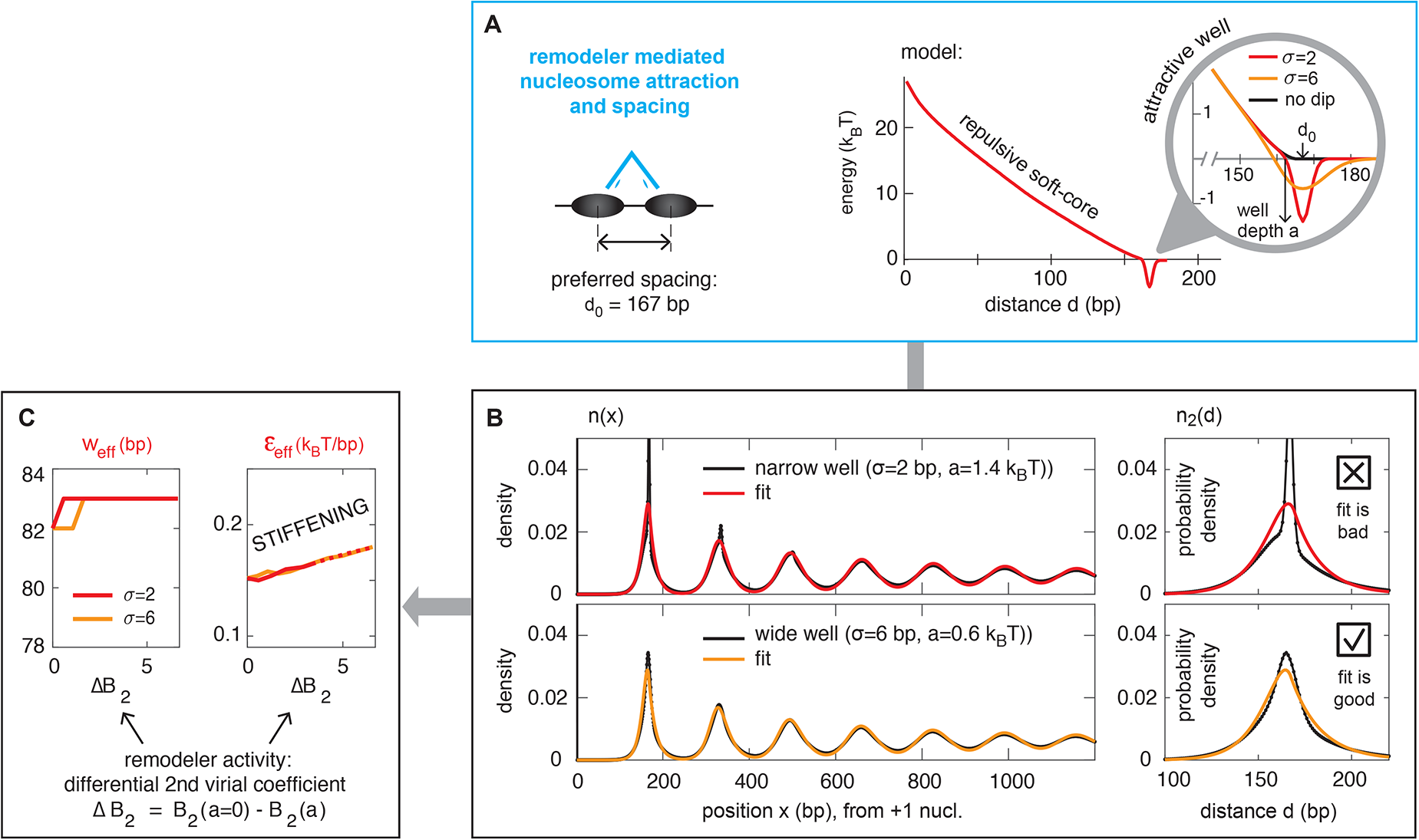
Effective nucleosome stiffening by remodeler mediated nucleosome attraction and spacing. (A) Attraction and spacing activity are modelled by superimposing the soft-core repulsion with an attractive potential well of width *σ*, depth *a* and center position *d*_0_. (B) For high spacing accuracy (*σ* = 2 bp, top panels) the density *n*(*x*) and especially the spacing distribution *n*_2_(*d*) show pronounced spikes at the preferred spacing *d*_0_ that are incompatible with a description by the nucleosome gas model. (C) The effective nucleosome properties show a stiffening with increasing remodeler activity (measured by the change in the second virial coefficient due to the attractive potential wells, see Supplement). Dashed lines indicate bad compatibility with the nucleosome gas model (Supplement).

We find that high nucleosome spacing accuracy (*σ* = 2 bp) results in a strong peak in the density *n*(*x*) and especially in the spacing distribution *n*_2_(*d*)(Fig. 5B, top panels, see Fig. S6 for a parameter sweep of *a*). Consequently, the compatibility with the nucleosome gas model, which lacks this spacing peak, is not good. For a more sloppy nucleosome spacing (*σ* = 6 bp), however, the compatibility is better (Fig. 5B, bottom panels). We find that for accurate and sloppy spacing alike the effective particle stiffness *ε*_eff_ increases with the remodeler activity (Fig. 5C). This is expected, since nucleosomes are preferentially kept at a fixed distance instead of invading each other. In conclusion, we found that a somewhat sloppy remodeler mediated nucleosome attraction towards a preferred spacing can be effectively described by the nucleosome gas model with increased nucleosome stiffness.

### *AT* **trapping explains unusually strong oscillations in the** *S. pombe* **nuclesome density**

We have shown that gene averaged nucleosome patterns can be modified by a multitude of mechanisms which potentially differ between species. This puts us in a position to ask why nucleosomes appear much stiffer in *S. pombe* than in *S. cerevisiae* (Fig. 1C,D), even though the underlying histones are conserved. We suggest that the large effective stiffness results from an *S. pombe* specific positioning mechanism, namely nucleosome trapping on AT-rich sequences. We motivate our hypothesis by the observation from Moyle-Heyrman et al. (Moyle-Heyrman et al., 2013) that in *S. pombe* nucleosomes are preferentially located on AT rich sequences within coding regions. This correlation is even more pronounced in our alignment of genes by their +1 nucleosome (Fig. 6A), but still completely absent in *S. cerevisiae* (Fig. 6B).

**FIG. 6:**
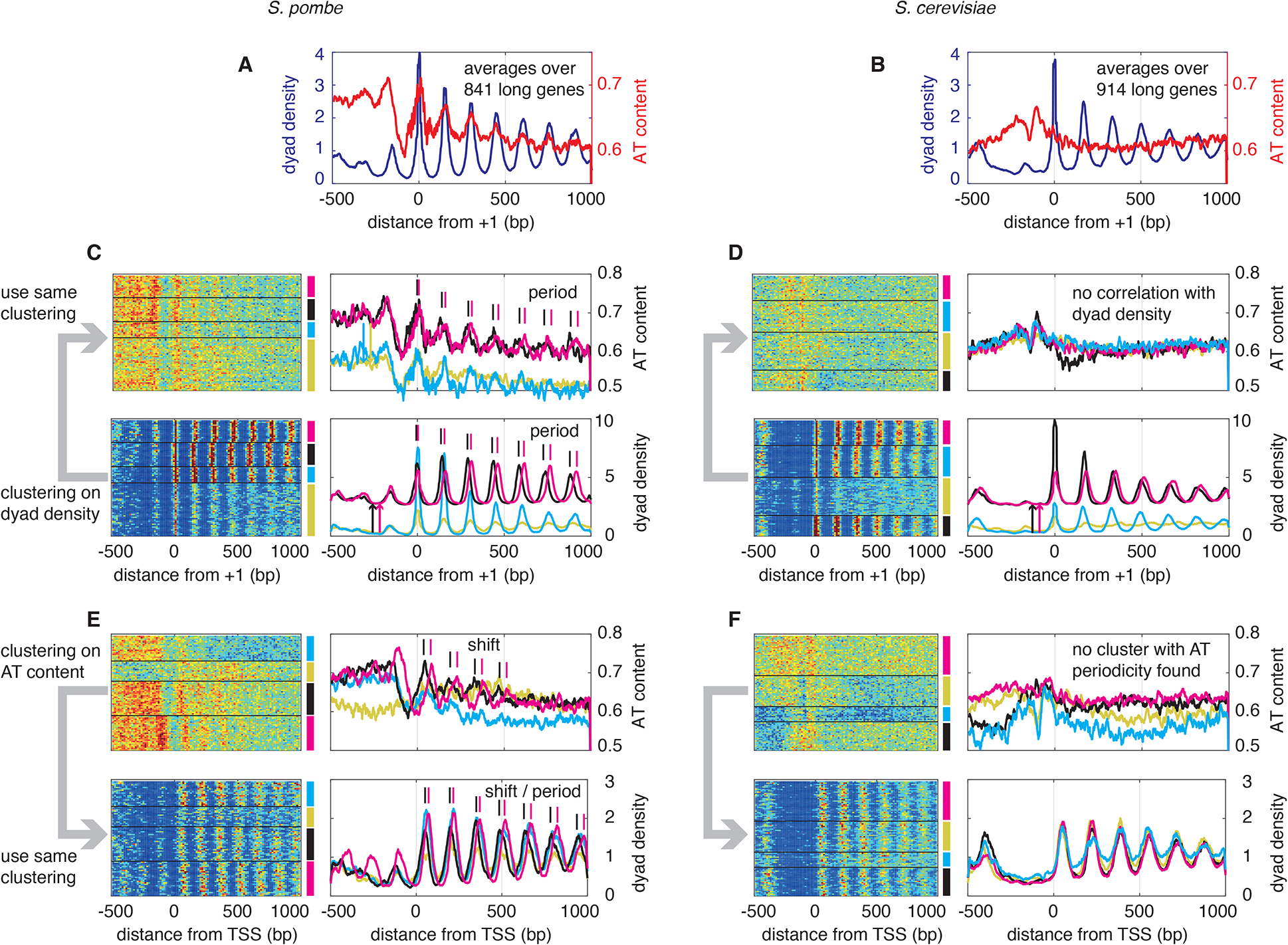
Nucleosome density and AT content strongly correlate in *S. pombe* but not in *S. cerevisiae*. Nucleosome density and AT content in *S. pombe* (A) but not in *S. cerevisiae* (B). This motivates our hypothesis that nucleosomes are trapped on AT rich sequences in *S. pombe*. Clustering genes based on their nucleosome dyad density (C,D, +1 aligned) or their AT content (E,F, TSS aligend) reveals additional correlations in *S. pombe* (see main text). Data are smoothed by 15 bp in x-direction. Dyad densities are normalized to unity on each gene. Some traces are offset for clarity as indicated by arrows.

To corroborate our hypothesis we dissect the correlations between nucleosome positions and AT content in more detail (Fig. 6C-F). We subdivide our set of long genes (> 2500 bp) in four clusters using *k*–means clustering (other numbers of clusters yield the same conclusions). First, we perform the clustering on the nucleosome density and check for correlations in AT content. We find that two clusters that differ in peak to peak distance (black and magenta lines in Fig. 6C), also show changes in the AT content. Furthermore, genes with weakly pronounced density peaks (pale green line) also show weak oscillations in the AT content. *S. cerevisiae* shows similar clusters in the nucleosome density but *none* of the features in the AT content (Fig. 6D). Next, we perform the clustering on the AT content and check correlations in nucleosome density. We find that in *S. pombe* shifted AT peaks are accompanied by shifted nucleosome patterns (black and magenta lines in Fig. 6E). In *S. cerevisiae* no genes with pronounced AT periodicity are found (Fig. 6F).

We conclude that nucleosome positions and AT content correlate strongly in *S. pombe* and we thus posit an energy landscape that preferentially positions nucleosomes on AT rich regions (Fig. 7B). Its period *λ* = 150 bp and decay length *l*_0_ = 555 bp are derived from the AT content oscillations (Fig. S7B).

**FIG. 7:**
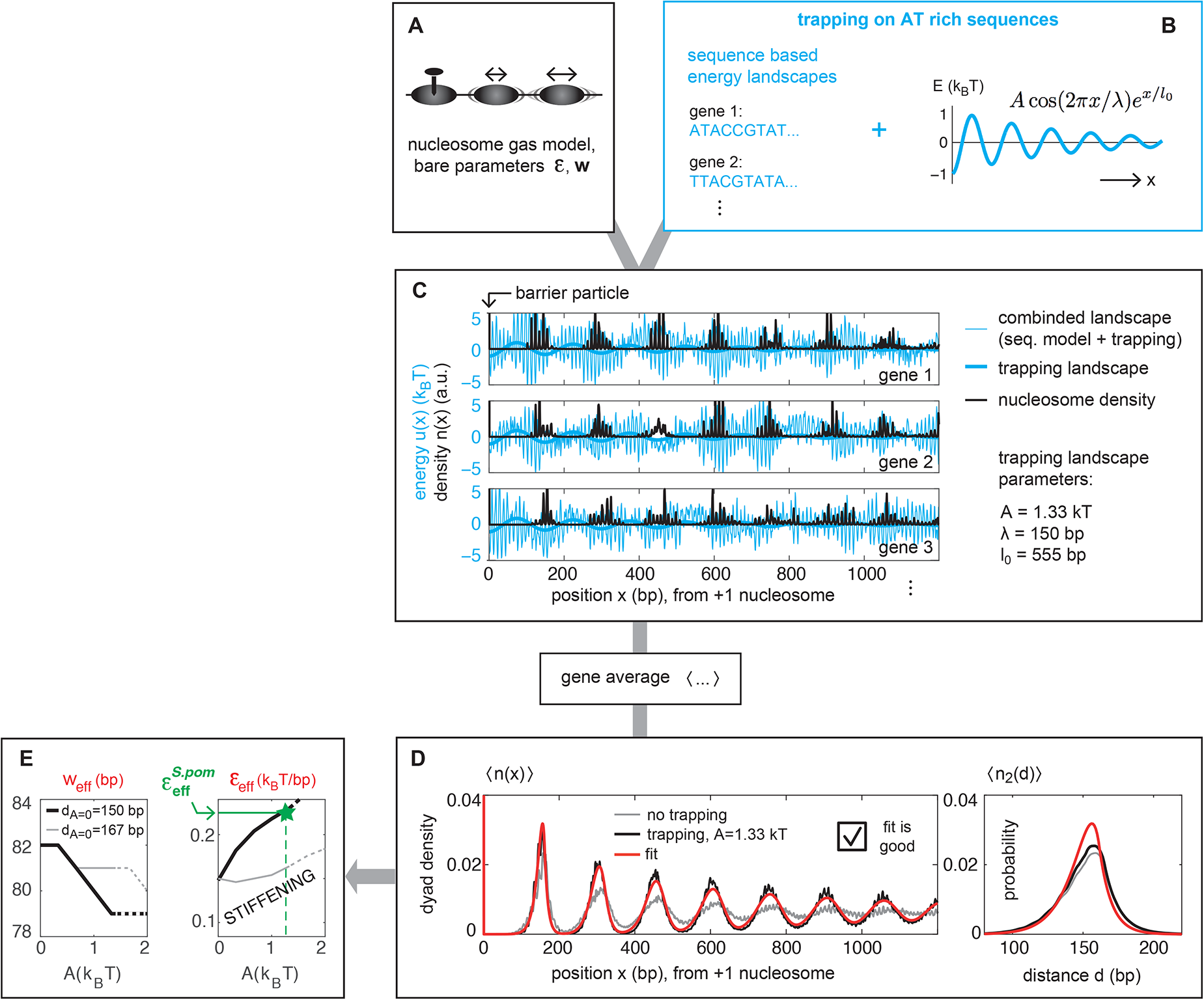
Effective nucleosome stiffening by trapping on AT rich sequences in *S. pombe.* Similar to Fig. 3 we use (A) the nucleosome gas model together with (B) DNA sequence specific energy landscapes, but now superimposed by a decaying oscillatory potential (thick cyan line) to account for trapping on AT-rich sequences as identified in Fig. 6. (C) Exemplary energy landscapes (cyan) together with the gene-specific nucleosome densities (black). (D) The gene averaged nucleosome density 〈*n*(*x*)〉 shows oscillations that are reinforced by the AT trapping (black *vs.* grey) and can be very well reproduced by the nucleosome gas model (red). (E) The effective nucleosome properties show a stiffening for in creased AT trapping strength (see text for details). Dashed lines indicate bad compatibility with the nucleosome gas model (Supplement).

We point out that the sequence based model for energy landscapes used above predicts no AT positioning on average on *S. pombe* genes (Fig. S3E). While it will be interesting to determine if other sequence based models do so, or if an indirect mechanism like remodeling has to be invoked, this question is irrelevant for our current goal to show that AT trapping can explain the large effective nucleosome stiffness in *S. pombe.* Here, we account for the combined effect of DNA sequence specificity and AT trapping by superimposing sequence derived landscapes and the trapping landscape. Examples are shown in Fig. 7B.

To account for AT trapping within the nucleosome gas model we first compute 〈*n*(*x*)〉 and 〈*n*_2_(*d*)〉 with AT trapping. On single genes the effect of AT trapping is rather weak (Fig. S7C), but when averaged over many genes the resulting oscillations are considerably strengthened (black vs grey line in Fig. 7D). Next we fit the gene averaged density and spacing distribution by the nucleosome gas model and determine the effective nucleosome parameters from the fit. We find that AT trapping renormalizes the nucleosome stiffness *ε* to larger values, *ε*_eff_ > *ε* (black line in Fig. 7E). This effective stiffening reaches *ε*_eff_ = 0.23 *k_B_T* (green star), which is close to the *S. pombe* value of *ε*_eff_ = 0.236 *k_B_T* (green solid line), at a trapping strength of *A* = 1.33 *k_B_T* (greed dashed line). This trapping strength is surprisingly low compared to the typical oscillations of sequence based landscapes (compare the bold and thin cyan lines in Fig. 7C) reflecting that nucleosomes guide each other by repelling reach other. The much slower decay of the density oscillations (compare black to grey line in Fig. 7D) is completely consistent with the much more pronounced oscillations in experimental *S. pombe* data compared to *S. cerevisiae* (Fig. 1C,D). We also observe that the effective nucleosome size parameter *w*_eff_ does not drop to the *S. pombe* value of 75 bp in the range suggested by the *ε* renormalization. We come back to this in the discussion.

We here assumed a nucleosome density whose oscillation period matches the *S. pombe* value of *d* = 150 bp even without the trapping landscape, the only effect of which was to result in more persistent oscillations in the gene body. For completeness, we also consider the case that the native density oscillation period equals the *S. cerevisiae* value of *d_A_*_=0_ = 167 bp. We find that cranking up the AT trapping imposes its period of 150 bp on the density and, for sufficiently strong trapping, again leads to effective nucleosome stiffening (thin grey line in Fig. 7E).

We conclude that nucleosome trapping on AT-rich sequences, which is motivated by strong correlations between nucleosome density and AT-content in *S. pombe*, can explain why nucleosomes appear much stiffer (i.e. positioning oscillations are more pronounced) in *S. pombe* than in *S. cerevisiae* even though the underlying histones are conserved.

## DISCUSSION

Towards a quantitative understanding of nucleosome positioning we here asked how different molecular mechanisms interact. We particularly focussed on how such mechanisms are reflected in gene-averaged positioning data. The essence of gene-averaged nucleosome patterns can be understood by the surprisingly simple model of statistical positioning against a barrier (Kornberg and Stryer, 1988; Mavrich et al., 2008). We showed that a refined version of statistical positioning, where DNA unwrapping is taken into account via a soft-core nucleosome interaction, accurately describes the characteristic oscillations in gene coding regions as well as the distribution of inter-nucleosomal spacings. This left us with an apparent contradiction: while *gene-specific* positioning preferences are not required to reproduce gene averaged patterns, such preferences are very clearly present and indeed shape gene-specific nucleosome patterns (Fig. 1B). We therefore investigated how gene-specific positioning alters gene-averaged patterns. We found that it changes the gene-averaged patterns quantitatively but not qualitatively. This was true independently of whether gene-specific positioning was generated from a DNA elasticity model or randomly, which indicates that the compatibility of strong positioning on individual genes and the emergence of simple nucleosome gas features in the gene average are a generic phenomenon.

Going beyond the qualitative finding that specific positioning mechanisms can be compatible with the nucleosome gas model in the gene average, we investigated their quantitative effects. Our general approach was to first model the nucleosome gas *together* with some specific positioning mechanism, and to then reproduce the obtained patterns with *only* the nucleosome gas model (Fig. 2). We found that the resulting effective nucleosome properties, in particular the effective energy required for unwrapping DNA from the histone core (the “nucleosome stiffness”), is renormalized by the additional positioning mechanisms. We would thus argue that Kornberg and Stryer’s original conclusion, that “the sequence-specificity cannot bee too great” (Kornberg and Stryer, 1988) should be modified to “the trace of sequence-specificity in the gene-average is a renormalization of the effective nucleosome properties”.

We found that different positioning mechanisms alter the effective nucleosome properties in different ways: While DNA sequence preferences (Fig. 3) and remodeler mediated nucleosome sliding (Fig. 4) lead to an effective softening, the setting of a preferred spacing (Fig. 5) or positioning along genes (Fig. 7) makes nucleosomes appear stiffer. We suggested that the latter can resolve the apparent conflict that DNA unwrapping seems particularly hard in *S. pombe* even though the underlying histones are conserved.

The picture that emerges from our results is that the salient feature of gene averaged nucleosome patterns, the decaying oscillations, emerges robustly from steric exclusion on DNA. The quantitative aspects of these oscillations, however, reflect the additional positioning mechanisms, which can be specific to species, cell types and conditions.

Our findings suggests that the quantitative features of nucleosome positioning data can be used to unravel the importance of specific positioning mechanisms. This requires some care, though. First, we have seen that some mechanisms lead to effective nucleosome stiffening and others to softening, and different contributions could thus cancel. Furthermore, different mechanisms can produce similar parameter renormalizations. For example, softening is caused by gene averaged DNA sequence effects as well as by remodeler mediated sliding, and stiffening could come from either a preferred nucleosome spacing (a two-particle quantity) or from preferred positioning along the genome (a single-particle quantity). With respect to the anomalously large unwrapping energy in *S. pombe*, which we suggested to stem from trapping on AT rich regions, alternative appealing scenarios are altered interactions due to the lack of the linker histone H1 or altered remodeling due to the lack of the ISWI remodeler in *S. pombe.* Further targeted studies are required to disentangle those effects.

In spite of the above mentioned cautionary notes our quantitative approach can greatly help in disentangling the interplay of positioning determinants. For example, we have demonstrated that the conjectured nucleosome spacing by remodelers must be either somewhat fuzzy, or can not be a dominant mechanism *in vivo* (Fig. 5). Furthermore, the correlation between AT content and nucleosome density in *S. pombe* suggested an AT trapping mechanism.

We point out that gene averaged patterns reveal information that would be hard to gain from single genes or specific loci, as illustrated by the above examples of AT trapping and fuzzy spacing. Of course, gene-specific patterns carry a great amount of additional information, which can be explored in the future. This will build on the here developed tools of quantitative simulation of specific remodeling mechanisms together with nucleosome interactions and sequence specificity.

Finally, we address experimental ramifications of our results. First, we suggest that the additional information revealed by chemical cleavage data over conventional MNase maps, namely the distribution of internucleosomal spacings (Fig. 1E), is extremely valuable: much more sensitive to mechanisms affecting the spacing than might be apparent from the nucleosome density patterns (Fig. 5B). Furthermore, it provides information that is simply not contained in the density patterns (Fig. 1E). In addition, the strength of oscillations in gene-averaged patterns depends on the pronouncedness of the barrier (no barrier, no oscillations). The barrier likely results from an interplay of sequence (Segal et al., 2006), remodeling (Krietenstein et al., 2016) and transcription (Chereji et al., 2016) and may vary across species and conditions. This again highlights the importance of the spacing distribution as a source of information about spacing mechanisms, since it does not rely on a barrier but rather reflects the nucleosome-nucleosome interaction directly. Measuring it could be of particular relevance in *in vitro* reconstitution experiments (Zhang et al., 2011), especially with purified remodelers (Krietenstein et al., 2016) in order to unravel their spacing activity.

Furthermore, trans-species experiments have proven useful: nucleosomes on *K. lactis* sequences cloned into *S. cerevisiae* were spaced with the shorter *S. cerevisiae* spacing, indicating that *trans* acting factors determined the spacing (Hughes et al., 2012). Similarly, replacing the endogenous gene for the remodeler Chd1 in *S. cerevisiae* by its *K. lactis* ortholog led to slightly increased nucleosome spacing (Hughes and Rando, 2016). Similar experiments with *S. pombe* might reveal the mechanism behind the correlation of nucleosome density and AT content.

In conclusion, we have established a framework for the quantitative understanding of how various mechanisms interact in nucleosome positioning. This approach can be particularly helpful in addressing a gap in current knowledge about chromatin remodeling enzymes: while single molecule experiments have revealed basic relocation steps (Flaus and Owen-Hughes, 2011; Mueller-Planitz et al., 2013) it is largely unclear how such relocations are combined to shape characteristic nucleosome positioning patterns. An iterative procedure of identifying candidate relocation mechanisms from single molecule experiments, exploring their single gene or genome wide effects in simulations as done here, and comparison to knockout or reconstitution experiments can unravel the building blocks of nucleosome positioning. Furthermore, an extremely interesting perspective is to unravel the relation between 1D nucleosome positioning and the 3D organization of chromatin. With Micro-C (Hsieh et al., 2015), which produces nucleosome positioning maps as well as 3D contact frequencies, questions about the 1D-3D relation might become addressable by joint experimental and modeling efforts.

## Acknowledgements

We acknowledge helpful discussion with Philipp Korber. MW is supported by SFB863 and a member of the Graduate School of Quantitative Biosciences Munich.

## Author contributions

Conceptualization: J.N. and U.G.; Investigation: J.N., M.W., B.O., W.M., and U.G.; Writing Original Draft: J.N. and U.G., with contributions from M.W., B.O., W.M.; Funding Acquisition: U.G.

## Supplement

### S1. THE SOFT-CORE NUCLEOSOME GAS MODEL

#### A. Nucleosome gas models

By a “nucleosome gas model” we understand the following model for nucleosomes on DNA:

- Nucleosomes are interacting particles on a discrete one-dimensional (1D) substrate, i.e. the DNA.
- The nucleosome-nucleosome interaction is a function of the separation *d* = |*x*_2_ − *x*_1_|, where *x*_1_ and *x*_2_ are the two nucleosome positions (measured by nucleosomal reference points, e.g. the nucleosome dyads):

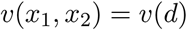

Thus, the nucleosome interaction does not depend on the position of the nucleosome pair, but only on the relative distance.
- There is no energetic preference for nucleosomes along the 1D substrate (note, however, that sometimes we introduce a barrier by a strong energetic preference at a single site, namely at *x* = 0).
- The above defined interacting lattice gas is in thermodynamic equilibrium.
- Depending on convenience, one can either use the *canonical ensemble* with a fixed number of particles or the *grand canonical ensemble* where particles can bind to and unbind from the 1D substrate, with a rate ratio defined by the chemical potential *r*_+_/*r*_−_ = *e^μ^*. Note that in large systems, which we are interested in, the two ensembles are equivalent.

Note that we defined the nucleosome gas model here to have no energetic preference along the DNA (i.e. no “energy landscape”). This is an arbitrary choice and due to the fact that we treat energy landscapes as additional positioning mechanisms.

#### B. Hard-core and soft-core interaction

The nucleosome-nucleosome interaction can be either a hard-core interaction, where each particle occupies a certain number of base pairs and overlaps between particle footprints are not allowed, or a soft-core interaction where overlaps are penalized by an interaction energy. The hard-core model is a special case of the soft-core model where the interaction energy tends to infinity when overlap occurs and zero otherwise. With the hard-core interaction, the model corresponds to the Tonks gas known in statistical physics (Tonks, 1936).

#### C. The soft-core interaction used in this paper

Here we use a previously introduced soft-core interaction (Möbius et al., 2013; Osberg et al., 2015, 2014) in order to account for nucleosomes invading each other on DNA (see below for interpretation). The soft-core interaction has two parameters, *ε* and *w*. The nucleosome stiffness parameter *ε* is the energetic cost of unwrapping one bp of DNA from the histone core. The nucleosome footprint parameter *w* determines the footprint of the fully wrapped nucleosome as 2*w* + 1 (see Fig. 1 in the main text).

The interaction potential for a given separation *d* between two nucleosome dyads is derived from summing the statistical weights of all unwrapping states of the left and right nucleosome that are compatible with the given separation *d* assuming thermodynamic equilibrium (see (Möbius et al., 2013) and Figs. 5A and S1D,E for graphs of the interaction potential).

#### D. Interpretation of the soft-core interaction potential

The soft-core interaction was motivated by unwrapping of DNA from histone core, but it can also represent other aspects of effective interaction between nucleosomes. Examples are steric hindrance of nucleosomes in 3D, effects of linker histones, and the presence of (active) remodeling enzymes. In particular the presence of H1 linker histones might contribute to increasing the effective footprint size beyond the 147 bp observed in nucleosome crystal structures. In order to take those aspects into account, we did not fix the parameters *ε* and *w*, but rather obtained them from fitting the model to experimental nucleosome positioning data (Fig. 1).

We note that the dyad-to-dyad spacing distributions of both *S. cerevisiae* and *S. pombe* show a finer substructure than our nucleosome gas model can reproduce: DNA unwrapping preferentially occurs in 10 bp steps due to “contact points” related to the helical twist (Luger et al., 1997). While a recent study has addressed this intranucleosomal structure (Chereji and Morozov, 2014), we here use a more coarse grained approach neglecting this fine structure.

#### E. Comparison of nucleosome interaction potentials from unwrapping model and as inferred from the experimental spacing distributions

Above, we have introduced a biologically motivated model for the soft-core nucleosome interaction based on DNA unwrapping, who’s parameters are determined by fitting of the model to data (Fig. 1 and below). For comparison, we here start with the experimental spacing distribution and reverse engineer the interaction, as based on a calculation from Ref. (Hansen-Goos et al., 2007). In brief, we use that spacings are exponentially suppressed by their length *d* and their energetic penalty *v*(*d*) such that the spacing probability density is

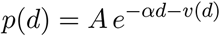

Thus, spacings decay exponentially for *d* large enough such that *v*(*d*) = 0. From this, *A* and *α* can be obtained by a linear fit to log(*p*(*d*)). Then, for smaller *d*, we obtain the interaction *v*(*d*) as follows:

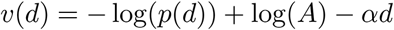

For the linear fit to determine *A* and *α* we used the spacing distribution in the range 170–180 bp. This is motivated by the fact that (i) for both species the fit of our unwrapping model to the data yields a vanishing interaction in the above range (for *S. cerevisiae* / *S. pombe* we have 2*w* + 1 = 165 / 151 bp as maximal interaction range, see Fig. 1), and (ii) the spacing distribution at increasingly long reads likely can not be trusted due to loss of long reads.

Results are shown in (Fig. S1D,E). We find very good agreement between the unwrapping model and reverse engineered potential.

### S2. COMPUTING THE NUCLEOSOME DENSITY AND SPACING DISTRIBUTION

As stated above, for the “nucleosome gas model” we assume no energetic preference for nucleosomes along the DNA. However, we later consider such energetic preferences (“energy landscapes”) as an additional positioning mechanism. Here, we thus use computational methods to obtain the nucleosome density and the spacing distribution in the presence of such an energy landscape, understanding that for the nucleosome gas model the landscape will be flat.

The problem of computing the one- and two-particle densities of particles with short-range interaction on a 1D lattice can be solved with tools from statistical mechanics. In particular, an arbitrary position dependent external potential, an “energy landscape” can be taken into account, as described in the works of J. Percus (Percus, 1989). We compute the one-particle density *n*(*x*) and the two-particle density *n*_2_(*d*) with a straightforward implementation of formulas in this reference.

Importantly, these methods assume that the system is in thermodynamic equilibrium. This is a strong and debatable assumption, given that active chromatin remodeling enzymes play an important role. Part of our results, however, show that the equilibrium-based soft core gas model indeed reproduces experimental data well (Fig. 1). Other parts of our results rationalize this agreement by showing that some remodeling mechanisms, e.g. nucleosome sliding (Fig. 4) are excellently compatible with an effective equilibrium description.

For the nucleosome density in gene-coding regions we assume that one particle, the “barrier particle” or “+1 nucleosome” is fixed in place and phases the adjacent particles. We fix this particle by placing a deep energetic minimum of −35 *k_B_T* at the desired position.

The relevant predictions of the nucleosome gas model are the density downstream of a barrier particle *n*^model^(*x*) and the nearest-neighbor spacing distribution 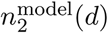.

When comparing the nucleosome gas quantities to experimental data we need to take into account that the reference nucleosome, which acts as a barrier particle, is not perfectly localized. We thus convolve *n*^model^(*x*) with the experimental +1 peak form 50 bp left to 50 bp right of the alignment point, resulting in *ñ*^model^(*x*). For simulated “data” this step is omitted.

### S3. FITTING THE NUCLEOSOME GAS MODEL TO DATA

We fit the prediction of the nucleosome gas model (see above) to data. Here, “data” refers to either experimental positioning data or to data from computational models including an “additional positioning mechanism”.

#### A. Fit parameters

When fitting the nucleosome gas model to data we vary all three model parameters, *ε, w*, and *μ* (see above). However, the quantities of interest are only the nucleosome properties *ε* and *w* and we report only those. The chemical potential *μ* is not an observable that we can compare to experimental data and we thus treat it as a dummy parameter.

To perform the fit, we choose a grid in the *w, ε*, 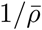 parameter space with a resolution of Δ*ω* = 1 bp, Δ*ε* = 0.002 *k_B_T* and 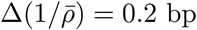. Here, 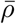 is the average nucleosome density and 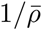 the average nucleosome spacing. We use this quantity to discretise the parameter space because the relation between the chemical potential *μ* and 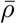 is highly non-linear.

#### B. Preparation of data for fitting by the nucleosome gas model

We use “chemical cleavage” nucleosome positioning data (Brogaard et al., 2012; Moyle-Heyrman et al., 2013). Instead of the Bayesian deconvolution technique used in Ref. (Brogaard et al., 2012), where the two nucleosome positions defining the endpoints of the read (read mates) were treated independently, we instead consider the mate pairs as a quasi-2D measure for nucleosome position and interdyad distance simultaneously, with some fuzziness or noise introduced by the cleavage bias as observed in Ref. (Brogaard et al., 2012). We then reconstruct the original 2D position-distance map using image deconvolution. This requires the specification of a point spread function (PSF). The PSF is calculated from the “single-end cleavage bias” (or “template weighted score” in Ref. (Brogaard et al., 2012)) by combining the effect of independent cleavage effects on position and distance. Adding 500bp flanking sequence around each gene, we then perform the deconvolution using the Matlab library function deconvlucy.

From the obtained genome wide nucleosome dyad density we use only genes longer than 2500 bps (to minimize influence from the transcription termination site). We align genes by the most likely position of their first nucleosome (the +1 nucleosome), which we determine as follows: From the TSS we look for the first downstream nucleosome position; if none is found within 300 bp we loop up to 50 bp upstream. If none is found, the gene is discarded (51 for *S. cerevisiae*, 2 for *S. pombe*). This results in 914 genes for *S. cerevisiae* and 841 for *S. pombe.* The distribution of dyad-to-dyad distances is taken only from the selected genes.

For experimental nucleosome maps, the data is not normalized and depends for example on the sequencing depth. We thus infer a normalization from comparison to the model. This can be done analytically, as described in Ref. (Möbius et al., 2013). Basically, we compute the normalization parameter *α*, which we refer to as the ‘sequencing depth’, as

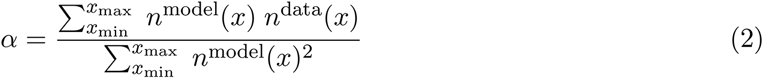

where *x*_min_ and *x*_max_ specify a valid range which we choose the same as the fit range (see below).

For the spacing distribution in experimental data the normalization is not known either, since reads beyond a certain length window [*d_min_*, *d_max_*] are lost. The nucleosome gas predictions, however, have tails beyond this window. Therefore, we equate the areas under the curves between the cutoffs. Namely, we use 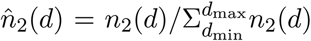 of the nucleosome gas prediction and the data. We use *d*_max_ = 200 bps as the upper cutoff when fitting experimental data and *d*_max_ = 300 when fitting simulated data, and we also use a sufficiently low *d*_min_ = 50 as lower cutoff.

As gene-averaged data may still be somewhat noisy we smooth the data prior to fitting by computing moving averages over 10 bp.

#### C. Fit error function

The quadratic error to be minimized is calculated as follows:

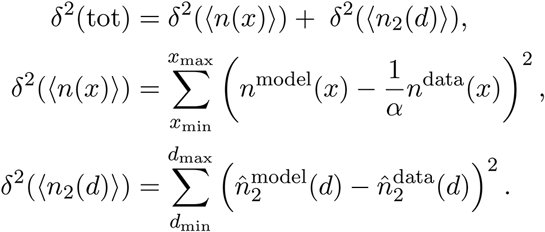

We choose the fit range *x*_min_ = 100 and *x*_max_ = 1500 for the density. The internucleosomal distances that are used for the fit of the gene- and position-averaged spacing distribution are taken only from this range as well. For the fit range for the spacing distribution we use [*d_min_*, *d_max_*] from above.

As the sum in *δ*^2^〈(*n*(*x*))〉 runs over more points than in *δ*^2^(〈*n_2_*(*d*)〉) the fit is more sensitive to the density. A possible way to adjust the relative weight would be to reduce *δ*^2^(〈*n*(*x*)〉) by the number of particles in the fit region 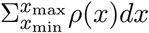. Yet, we find that this leads to very unsatisfying results for the density and thus refrain from the correction. We point out that, in any case, discrepancies point to deviations from Tonks gas physics, irrespective of whether the density fit is ‘improved’ at the cost of the spacing distribution or the other way round.

#### D. Criterion for god or bad fit

To decide if (experimental or simulated) data is compatible with the nucleosome gas model we need a criterion for what constitutes a good fit and a bad fit. Defining such a criterion is somewhat arbitrary. Here we choose a maximal value of the total fit error, namely *δ*^2^ (tot) < 0.003 for a good fit (magenta line in Figs. S2, S5, S6, S7.

We point out that the total fit error should also be compared to the effect of the additional positioning mechanism under consideration: ideally, the effect of the mechanism on the pattern is large, but the compatibility is still good (i.e. *δ*^2^(tot) is small). Thus, we plot also the squared deviation between our quantities with additional positioning mechanism and without the mechanism (“to unperturbed” in Figs. S2, S5, S6, S7).

### S4. FIT TO DATA FROM *S. CEREVISIAE* AND *S. POMBE*

In Fig. S1 we show the fit of the nucleosome gas model to the experimental data from *S. cerevisiae* and *S. pombe* (see Fig. 1 in the main text) alongside the density and spacing distribution from non-optimal parameters (purple and green lines).

We note that the fit error for the density alone, *δ*^2^(〈*n*(*x*)〉), has an extended banana-shaped valley with a comparatively sloppy mode. Consequently, the density can be reasonably well reproduced by values (*ε, w*) that are far from the optimal ones, provided that they lie in the valley (see purple lines in Fig. S1B,C for examples). The spacing distribution, however, is more sensitive and its fit error *δ*^2^(〈*n*_2_(*d*)〉) has less of a valley. Correspondingly, the spacing distribution for the purple lines in Fig. S1 is not a satisfactory fit.

#### A. Note on the discrepancy between nucleosome density and spacing distribution in *S. cerevisiae*

An intriguing feature of the chemical cleavage data from *S. cerevisiae* is the difference between the peak to peak distance in the density pattern of 167 bp and the mean length of fragments (collected only in the region of this pattern) of only 157 bp, which was pointed out before (Chereji and Morozov, 2014). We can see two possible explanations: (1) An experimental bias towards short fragments (note that the simultaneous fit fails in particular at reads exceeding 170 bp). (2) Mechanisms beyond statistical positioning, inducing more correlations.

### S5. NUCLEOSOME POSITIONING BY ENERGY LANDSCAPE MODELS

#### A. Landscape models

We use two types of energy landscapes: sequence based energy landscapes and random landscapes without an underlying physical model. We point out that our focus is not the predictive power of specific models at specific loci, but rather the generic effect of energy landscapes on gene-averaged patterns. We show that in this respect the details of the model are of minor importance.

As input for a sequence based landscape model we use *S. cerevisiae* and *S. pombe* DNA sequences aligned by the +1 nucleosome. We use only genes longer than 2500 bp in order to minimize effects from the transcription termination site, see above. We compute landscapes using 4000 bp downstream of TSSs. For the fit to the nucleosome gas, however, we use only the first 1500 bp as explained above. The large overhang is to avoid effects from the end of the substrate.

As a specific sequence based model for energy landscapes we use the model proposed in Ref. (Morozov et al., 2009). This computes the energetic cost of bending a specific DNA sequence into the shape required fro wrapping around the histone core based on elastic properties of the DNA. Namely, each base pair stacking is associated with deformation force constants (3 displacements, 3 rotation angles).

The authors of Ref. (Morozov et al., 2009) use either only the central 71 bp or the full 147 bp to compute nucleosome formation energies. The former is motivated by the central part binding DNA first and thereby already determining the nucleosome positions, while the later wrapping of the outer parts has no influence on the nucleosomes sequence preference. We here use the central 71 bps. We checked, however, that using the full 147 bp yields qualitatively similar results in terms of the here studied parameter renormalization by energy landscapes.

To investigate the overall effect of the landscape amplitude (measured by the standard deviation *σ*) we rescale the landscapes and determine the effective nucleosome properties as a function of *σ*.

#### B. Computing the nucleosome density and spacing distributions on energy landscapes

We compute the nucleosome density and spacing distribution on individual energy landscapes exactly. We use the transfer matrix based equilibrium methods from statistical mechanics as explained above.

For each landscape we first set the mean of this landscape to zero and then subtract the chemical potential *μ*. When fitting the model to data, the chemical potential is a fit parameter as described below. When generating data (e.g. the nucleosome density on sequence based energy landscapes considered in this section) the chemical potential is obtained by an iterative process such that, in the gene average, a specified average nucleosome density is reached.

#### C. Gene averaged energy landscapes show now positioning featues

We pointed out in the main text that it is crucial to take sequence dependent positioning energy landscapes *u*(*x*) into account for each gene separately and *then* average the obtained densities across genes:

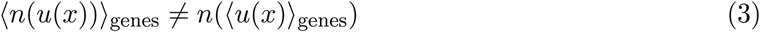

where 〈 〉_genes_ is the average across genes. For comparison, Fig. S3E shows the gene averaged energy landscape 〈*u*(*x*)〉_genes_. We observe that its standard deviation is very small, namely *σ* = 0.056 *k_B_T*, which is very close to 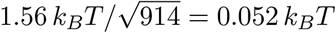, which is expected from averaging uncorrelated landscapes across our set of 914 genes. This indicates that there are no typical features encoded in the sequence dependent landscapes according to the used landscape model. These findings are independent of aligning genes by the +1 nucleosome or the TSS (Fig. S3E top and middle panel).

For energy landscapes from *S. pombe* sequences we observe more pronounced oscillations near the +1 nucleosome, which, however, disappear about 200 bp downstream, i.e. between the +2 and +3 nucleosome (Fig. S3E bottom panel). In particular, we observe no positioning oscillations in phase with the *S. pombe* specific AT content oscillations.

#### D. Detailed account of fitting of the nucleosome gas model to positioning data on energy landscapes

Figure S2 shows in detail the fitting of the nucleosome gas model to data from sequence based energy landscapes, corresponding to Fig. 3 in the main text. In order to determine if the data can be reproduced by the nucleosome gas model, we investigate the fit errors, see, panels (B). We show the error of the gene averaged quantities for a given *σ* with respect to the fit (red) and with respect to the quantities in the absence of landscapes (black). Thereby we compare the *compatibility* with the nucleosome gas model to the *effect* of energy landscapes on 〈*n*(*x*)〉 and 〈*n*_2_(*d*)〉. We find that the compatibility is very good compared to the effect (errors “to fit” are much smaller than error “to *σ* = 0”). For the error in the spacing distribution the two errors are comparable, which results from the fit being more sensitive to the density (see above). The magenta line in the total error plots indicates our cutoff for a good fit, namely *δ*^2^ = 0.001. All considered examples satisfy this criterion.

#### E. Inferring the amplitude of disorder landscapes from the peakedness of gene-specific nucleosome density profiles

In the main text we showed how gene specific nucleosome positioning energy landscapes make nucleosomes appear softened in gene-averaged patterns. We found that the degree of this parameter renormalization depends on the standard deviation of the energy landscapes. Here, our goal is to infer this standard deviation from the experimental nucleosome density profiles.

A natural observable to this end is the peakedness of the density on individual genes. Figure S3A shows the experimental *S. cerevisiae* dyad density for an exemplary gene, compared to model densities on a landscape with varying standard deviation *σ*. Importantly, all three cases are from the “intersection line” *ε* = 0.152*k_B_T* in Fig. 3A, namely, they all yield the same gene averaged patterns. On the single gene level, however, there are visible differences in the peakedness.

To quantify the peakedness, we propose histograms of the position dependent density values as an adequately integrated measure (Fig. S3B). For example, *σ* = 1.07 *k_B_T* leads to moderately peaked density patterns, where loci with large density peaks are less likely than in experimental data (the histogram drops much quicker than for the experimental data). Conversely, for *σ* = 2.48 *k_B_T* the model densities are too spiky and the corresponding histogram shows more counts for very large densities than the experimental data. The best agreement is obtained for *σ* = 1.56 *k_B_T*. This value is very consistent with the *σ* that the authors of the sequence based landscape model give, namely *σ* = 1.74 *k_B_T* (Morozov et al., 2009).

#### F. Parameter renormalization flow for sequence based energy landscapes and uncorrelated Gaussian landscapes

We pointed out in the main text that the main parameter for the renormalization of the nucleosome stiffness parameter *ε* is the standard deviation *σ* of energy landscapes. Here, we show that indeed a completely different landscape model leads to a very similar renormalization.

We use landscapes whose only parameter is the standard deviation *σ*. Landscapes are generated by drawing a random number from a Gaussian with standard deviation *σ*, 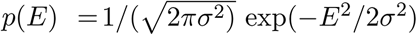, at each base pair independently (“uncorrelated Gaussian landscapes”). Example landscapes in Fig. S3C show that the typical 10 bp oscillation observed in sequence based landscapes (Fig. 3C) is absent here. Nevertheless, the corresponding parameter flow (Fig. S3D) is very similar.

#### G. Examples of positioning landscapes that are not compatible with the nucleosome gas model

In order to show that not all nucleosome density patterns near a barrier can be fitted, we tested two examples of random landscape ensembles that have a flat gene average, but result in densities that still cannot be fitted by the “interaction + barrier” model:

In the first example the external potential has normally distributed landscape values at each position with a non-constant standard deviation: introducing a peak in the standard deviation (max at 600bp, footprint 20bp, linear increase of multiplication factor from 1 to 11 and corresponding decrease) leads to potentially very high negative peaks in the landscape which strongly attract nucleosomes leading to a peak at 600bp in the one-particle density not in agreement with the normal nucleosome array pattern.

In the second example we artificially introduce correlations into the external potential. Starting from the barrier and moving downstream, negative peaks below −2.8 kT are copied further downstream by 172, 173, 174, 345, 346 and 347 base pairs, overwritting the original values. This step is repeated from the position of the last overwritten value until we reach the end of the region. This introduces correlations between strong negative peaks and leads to a bimodal spacing distribution with a second peak at 173bp without affecting the one-particle density. Strictly speaking this does not have a zero landscape on average yet, but this can be fixed by also copying peaks above +2.8 kT, without noticable effects on the average densities, since nucleosome positions are only governed by the lowest external potential values.

### S6. REMODELER MEDIATED NUCLEOSOME SLIDING

#### A. Model

In order to model nucleosomes subject to the action of active remodeling enzymes we use kinetic Monte Carlo simulations, which we describe in detail in this section.

We use a lattice of length *L* = 5010 with one lattice site corresponding to one base pair.

##### Nucleosomes

Nucleosomes can adsorb/desorb onto/from the lattice. We label nucleosomes by the position *x* of their center (their dyad) on the substrate. We use the following adsorption and desorption rates:

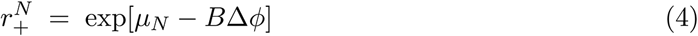

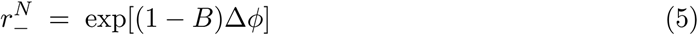

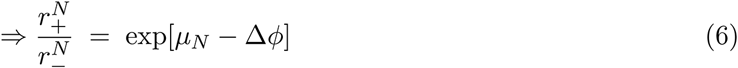

where *μ_N_* is the chemical potential for nucleosomes. *B* is a number between zero and one which *B* distributes the Boltzmann weight exp[Δ*ϕ*] to the rates of adsorption and desorption and thereby does affect the kinetics of each process separately, but not the rate ratio (see above) and thus not the relative equilibrium occupancies. In lack of detailed experimental data we use *B* = 1/2 throughout. See below for the particle-particle interaction potential change Δ*ϕ*.

##### Remodelers: adsorption-desorption

Remodelers can adsorb onto DNA bound nucleosomes. We use the rates

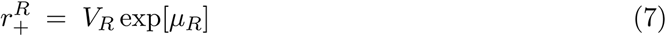

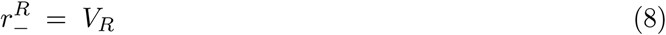

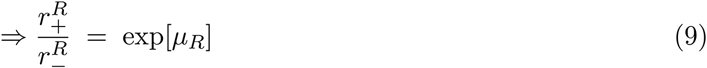

where *V_R_* is a basal “volatility” which sets the time scale of remodeler kinetics relative to the nucleosome kinetics, and *μ_R_* is the remodeler chemical potential. These considerations apply separately to every type of remodeler in our system (here, upstream and downstream remodelers, see below)

##### Remodeler-nucleosome complexes: adsorption-desorption

For consistency, we assume that remodeler-nucleosome complexes can also bind and unbind. We impose detailed balance for the reaction sequence 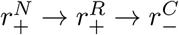, which means that *μ_C_* = *μ_Ν_* + *μ_R_*. We use the rates

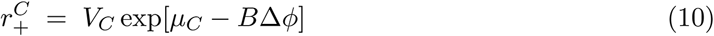

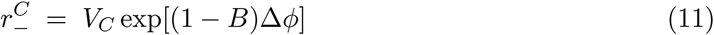

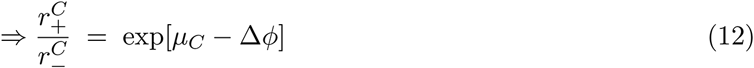

where *V_C_* sets the time scale of complex kinetics relative to the nucleosome kinetics. These considerations apply separately to every type of remodeler in our system (here, upstream and downstream remodelers, see below)

##### Interactions

All rates given above include modifications by particle-particle interactions which satisfy detailed balance. We assume that nucleosomes interact via the potential *ϕ* described above (see also (Möbius et al., 2013)). For simplicity we here also assume that nucleosome interactions are not modified by the presence of remodelers. This is likely not a realistic scenario, since (i) nucleosome remodelers are large proteins, and (ii) they can destabilise nucleosomes even in the absence of ATP [ref]. Yet, lacking a well-founded model for the interaction potential between remodelers, we here go forward with this simplifying assumption.

Rates are modified by

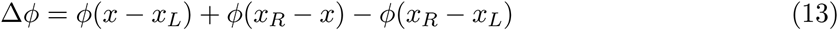

which is the change in interaction energy upon adsorption of a nucleosome or complex, where *x* is the center position of the adsorbing particle and *x_L_* and *x_R_* are the positions of the next particles to the left and right.

##### Remodeler mediated nucleosome sliding

As a minimal model for directional nucleosome sliding we consider a biased random walk: While bound to the nucleosome, an upstream/donwstream remodeler slides a nucleosome in a direction that is chosen at random, but with a bias for the upstream/downstream direction. For simplicity, we assume a stepsize of 1 bp. The sliding rates are

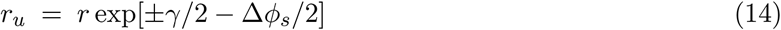

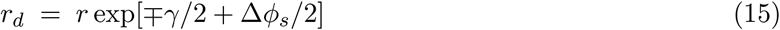

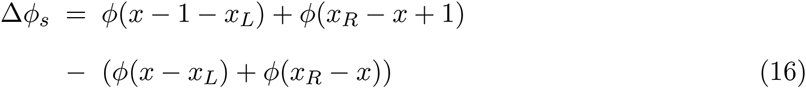

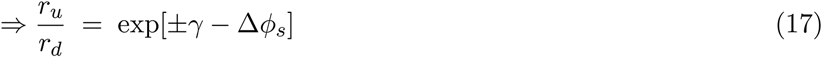

where *r_u_* and *r_d_* are the rates for sliding the nucleosome upstream from *x* and sliding the nucleosome downstream from the upstream position *x* + 1, respectivly. *r* is a rate (“basal activity”), which sets the timescale of remodeler mediated nucleosome sliding relative to the nucleosome adsorption/desorption kinetics, and the upper/lower sign is for upstream/downstream remodelers. Furthermore, if a sliding complex and another particle invade each other, the change in nucleosome interactions due to sliding, Δ*ϕ_s_*, is non-zero and slows down sliding reactions which increase the interaction energy, and expedites sliding which reduces the interaction energy by the same factor [Δ*ϕ_s_*/2].

The rate ratio (17) shows that, in the absence of interactions, an upstream/downstream remodeler preferentially slides a nucleosome upstream/downstream by a factor *e^γ^*. This sliding model breaks detailed balance and results in different steady state densities compared to a pure nucleosome adsorption/desorption model without sliding, if and only if *γ* ≠ 0.

##### Remodeler processivity

In order to measure the effectiveness of nucleosome sliding we define a remodeler’s processivity as the average displacement of a nucleosome during the residence time of the remodeler in the absence of any interactions. The processivity can be computed as follows: The average drift speed of the remodeler mediated random walk is the difference of downstream and upstream sliding rates, namely

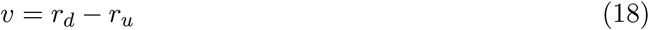

where we have chosen the downstream direction as positive. To obtain the average displacement we divide the drift speed by the average residence time, 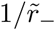, where 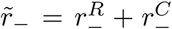 is the sum of the unbinding rates of the remodeler and the remodeler-nucleosome complex in the absence of interactions. We thus arrive at

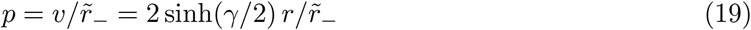

where for small *γ* we have

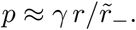

##### Simulation parameters for nucleosome sliding

We vary the remodeler processivity in two ways, namely by sweeping the bias parameter *γ* and the rate parameter *r*. We choose processivities that are of the order of *w*/2, thus of a quarter of the nucleosome size. Note, however, that processivities are defined in the absence of interactions. The actual remodeler mediated invasion of one nucleosome into another is smaller, since increasing interactions promote unbinding of the nucleosome-remodeler complex and furthermore progressively slow down the invasion and eventually neutralise and reverse the bias. The condition for neutralising the sliding bias due to another invaded particle is ±*γ* = Δ*ϕ_s_*. The used parameter values and resulting processivities are given in Table S1.

### B. Detailed account of fitting of the nucleosome gas model to remodeler mediated nucleosome sliding data

Figure S5 shows in detail the fits of the nucleosome gas model to the data from our model for remodeler mediated nucleosome sliding. The bottom graphs in panels A,B show the fit errors (red) and the deviation of the data in the presence of remodeling from data in the absence of remodeling (black). This juxtaposes the *compatibility* of the considered remodeling mechanism with the nucleosome gas model to the *effect* of this remodeling.

We find that the compatibility error is extremely low compared to the effect of nucleosome sliding. This indicates excellent compatibility of nucleosome sliding with the nucleosome gas model. The magenta line in the total error plots indicates our cutoff for a “good fit”, namely *δ*^2^ = 0.001. All considered examples satisfy this criterion.

### S7. REMODELER MEDIATED ATTRACTION AND SPACING

#### A. Model

As a coarse grained model for remodeler mediated nucleosome attraction and spacing we modify the repulsive soft-core nucleosome interaction by adding an attractive potential well:

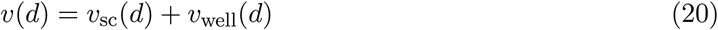

with

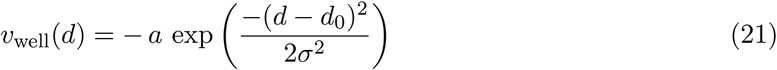

where *a* is the depth of the well (in units of *k_B_T*), *σ* the standard deviation and *d*_0_ the preferred nucleosome distance.

For our coarse grained model we compute the equilibrium density and spacing distribution with the above nucleosome interaction potential using the transfer matrix method as described above. We thus choose an equilibrium description of the effect achieved by energy consuming molecular machines, potentially neglecting any affects that are not amenable to such an effective description.

Within our coarse grained equilibrium model we sweep the remodelers’ attraction strength by changing the well parameter *a*. According to equilibrium statistical physics the correct measure for the impact of the attraction is the second virial coefficient, or, up to a factor −1/2, the integrated deviation of the Boltzmann factor from unity. In a 1D system it reads

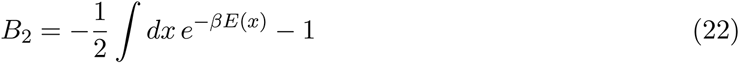

where *β* = 1/*k_B_T*. The integrand is shown in Fig. S6C. Contributions to *B*_2_ from repulsion are positive and contributions from attraction are negative. Due to the strong repulsive core our *B*_2_ are always positive (dominated by repulsion). We measure the impact of adding an attractive well to our soft-core repulsion by the corresponding change in the second virial coefficient as

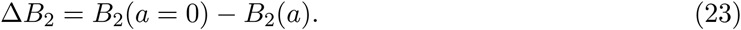

#### B. Detailed account of fitting the nucleosome gas model to remodeler mediated attraction and spacing data

Figure S6 shows in detail the fits of the nucleosome gas model to the data from our model for remodeler mediated attraction and spacing. The bottom graphs in panels A,B show the fit errors (red) and the deviation of the data in the presence of remodeling from data in the absence of remodeling (black). This juxtaposes the *compatibility* of the considered remodeling mechanism with the nucleosome gas model to the *effect* of this remodeling.

We find that for narrow wells (*σ* = 2) the compatibility error and the effect are similar, while for the wider wells (σ = 6) the compatibility error is much smaller than the effect, indicating better compatibility. The magenta line in the total error plots indicates our cutoff for a “good fit”, namely δ^2^ = 0.001. For the narrow well only low spacing activity (*a* ≤ 0.6) satisfies this criterion.

### S8. AT TRAPPING IN *S. POMBE*

#### A. Analysis of decaying oscillations of the AT content in +1 aligned *S. pombe* genes

Here we characterise the AT content in +1 aligned *S. pombe* genes. Fig. S7B shows the raw data for our set of 841 *S. pombe* genes longer than 2500 bp (grey), and the same data smoothed by a 30 pb sliding window. The latter clearly highlights out the oscillations.

Next we detect local maxima and minima (red and blue circles) in the smoothed data. Plotting the max-to-min difference for each consecutive pair on a log scale vs the position *x* of the maxima reveals an exponential decay of the max-to-min difference with *x*. The decay length is 1/0.001803 = 555 bp, the y-intercept is *2A* = exp(−2.686) = 0.068. The oscillation period λ as obtained from the maxima positions is 152 bp and matches excellently with the period of nucleosome density oscillations of 150 bp (Fig. 1D). For the AT trapping sequence we thus use

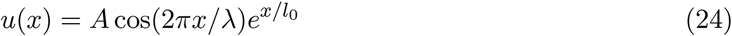

with λ = 150 bp and *l*_0_ = 555 and a variable *A* to investigate the nucleosome parameter renormalization for varying AT trapping strength. The corresponding relation is overlayed in green on panel Fig. S7B with the *A* = 0.034 from the fit (the offset and linear dependence are hand tuned to approximate the underground on top of which the maxima and minima are found).

#### B. Note on the interpretation of the AT trapping model in *S. pombe*

We have suggested that trapping on AT-rich sequences might explain why nucleosomes appear much stiffer in gene-averaged data from *S. pombe* due to parameter renormalization. We have not addressed what could cause this trapping. We have no indication that DNA bending mechanics alone is involved: The corresponding sequence based model (Morozov et al., 2009), when fed with the *S. pombe* sequences that exhibit 150 bp periodic AT content, does not show any landscape periodicity on average Fig. S3. An indirect effect, like sequence dependent remodeling, thus seems more likely. In any case, this distinction is unimportant for our statement that trapping makes nucleosomes appear stiffer.

The AT trapping mechanism renormalizes the footprint parameter *w* more than all other studied mechanisms. However, a renormalization to the value of 75 bp obtained from the fit of the soft-core model to the data seems out of reach for reasonable *ε* renormalization. It might be possible to obtain a stronger *w* change by tuning the shape of the trapping landscape.

**TABLE S1:** Simulation parameters for nucleosome sliding model. The lower two blocks are values used in the sweeps of the remodeler bias parameter and the rate parameter, respectively. Sweep values are given as a list in brackets. The resulting remodeler processivities are given as well. The nucleosome properties are chosen from the fit to *S. cerevisiae* (Fig. 1), namely *ε* = 0.152 *k_B_T* and *w* = 82 bp.

| parameter | description | value |
| --- | --- | --- |
| $\mu_N$ | nucleosome chemical potential | 4.012 |
| $V_R$ | remodeler timescale factor | 10 |
| $\mu_R$ | remodeler chemical potential | $-\ln(2)$ |
| $V_C$ | complex timescale factor | 1 |
| $\mu_C$ | complex chemical potential | $4.012 - \ln(2)$ |
| $B$ | Boltzmann weight distribution parameter | 1/2 |
| $r$ | rate parameter | $\frac{w}{2} \frac{V_R}{8\varepsilon}$ |
| $\gamma$ | bias parameter | $8\varepsilon[0, \frac{1}{2^4}, \frac{1}{2^3}, \frac{1}{2^2}, \frac{1}{2}, 1]$ |
| $p$ | processivity | $[0, 2.3, 4.7, 9.4, 19, 40]$ |
| $r$ | rate parameter | $\frac{w}{2} \frac{V_R}{\varepsilon} [\frac{1}{2^5}, \frac{1}{2^4}, \frac{1}{2^3}, \frac{1}{2^2}, \frac{1}{2}, 1, 2]$ |
| $\gamma$ | bias parameter | $\varepsilon$ |
| $p$ | processivity | $[1.2, 2.3, 4.7, 9.3, 19, 37, 75]$ |

**FIG. S1:**
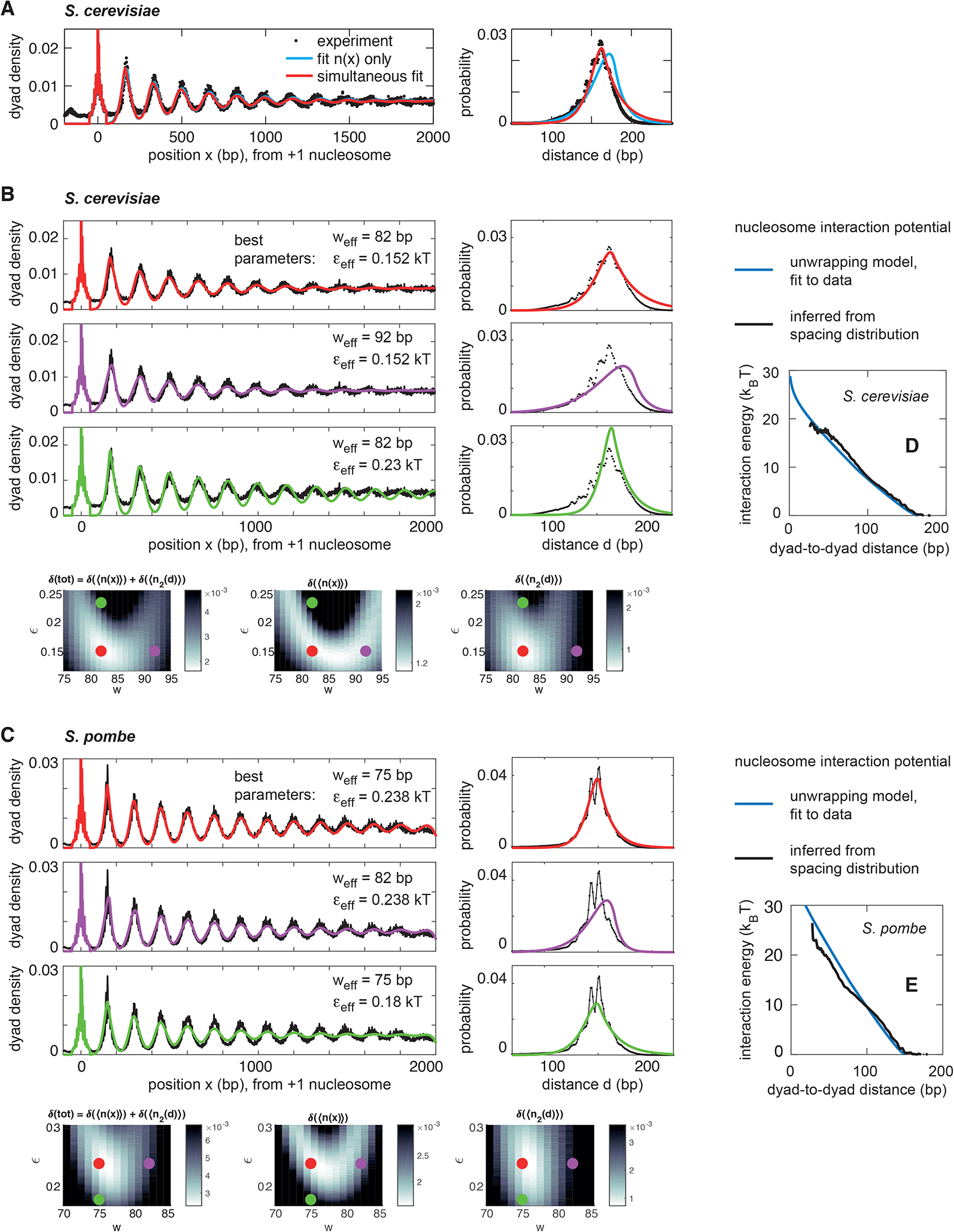
Related to Fig. 1. Fitting the nucleosome gas model to data from *S. cerevisiae* and *S. pombe*. (A) Comparison of fitting the nucleosome gas model to either the nucleosome density alone (blue) or simultaneously to the nucleosome density and the spacing distribution (red). The latter leads to much better agreement for the spacing distribution. This indicates two things: first, the density alone does not constrain the fit (see panels B,C). Second, there is some discrepancy between the density 〈*n*(*x*)〉 and the spacing distribution 〈*n*_2_(*d*)〉, since otherwise the fit to one of them should also reproduce the best fit to the other. At the moment we can only speculate that this discrepancy is due to the length selection of sequenced DNA fragments. We point out that this discrepancy is almost absent in *S. pombe* (Fig. 1D), which has a narrower fragment length distribution and might thus be less susceptible to length cutoffs. (B,C) Best fit (red) compared to other choices of *w* (purple) or *ε* (green). Again, non-optimal parameters impair consistency with the spacing distribution visually more than with the density. (D,E) Soft-core nucleosome interaction potential according to our model based on DNA unwrapping and fit to experimental data, compared to the potential inferred from the dyad-to-dyad spacing distributions.

**FIG. S2:**
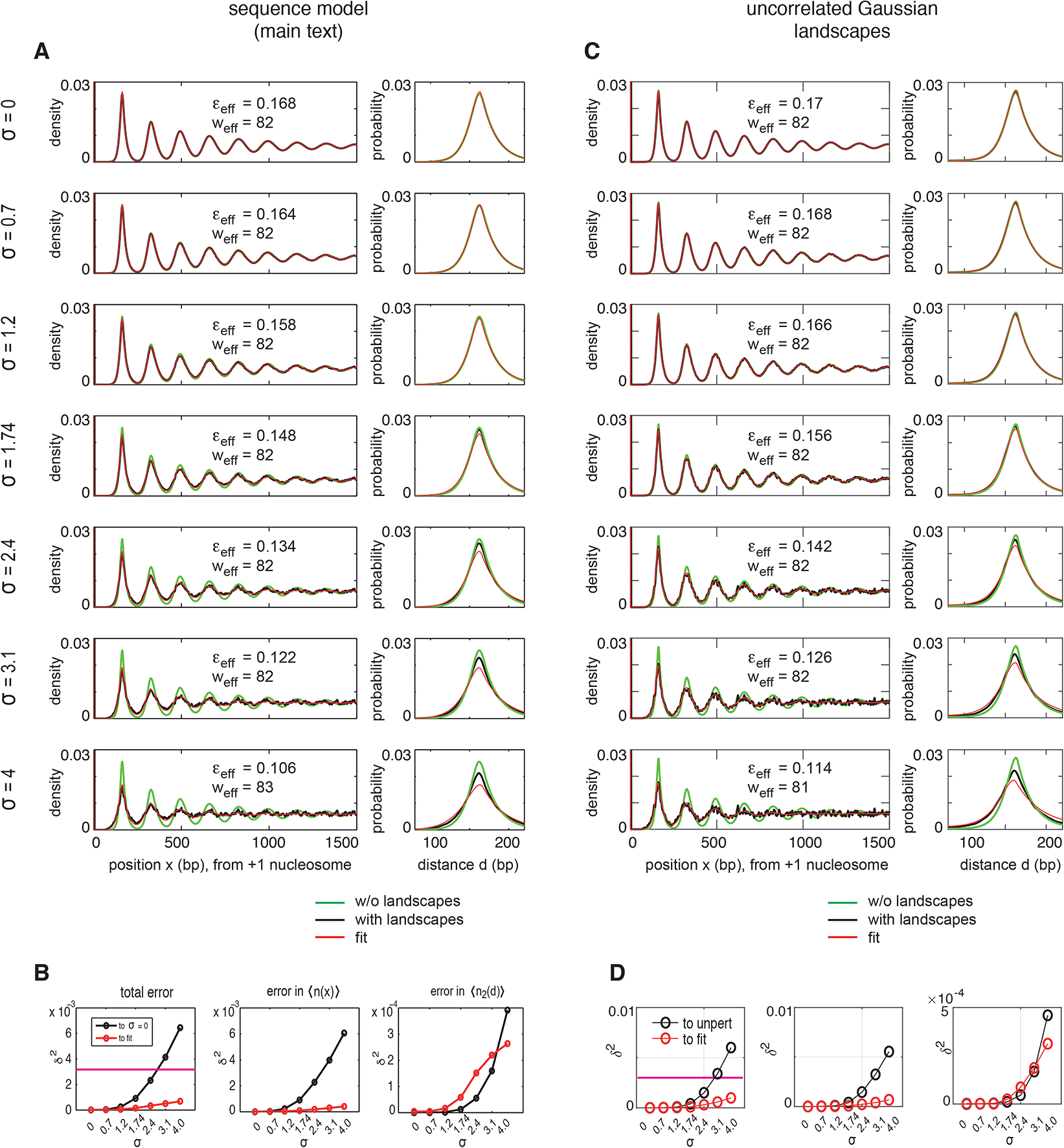
Related to Fig. 3. Fitting the nucleosome gas model to data from nucleosomes on energy landscapes. Ensemble averages of the nucleosome density 〈*n*(*x*)〉 (left panels) and spacing distribution 〈*n*_2_(*d*)〉 (right panels) on energy landscapes (black, smoothed by 10 bp). (A) Landscapes are computed from the sequence based DNA elasticity model (Morozov et al., 2009) using the DNA sequences of 914 *S. cerevisiae* genes that are longer than 2500 bp and aligned by the consensus position of the +1 nucleosome. (C) Uncorrelated Gaussian landscapes: on every gene at every basepair we draw the energy from a Gaussian with standard deviation *σ* to obtain uncorrelated landscapes. Examples are shown in Fig. S3C. (A,C) The quantities *n*(*x*) and *n*_2_(*d*) are computed on each gene as described above, using bare nucleosome parameters *ε* = 0.17*k_B_T*/bp and *w* = 82bp, and then averaged across all genes, yielding the black lines. The results are fitted by the nucleosome gas model without landscapes (red lines). The standard deviation *σ* of the landscapes is swept from 0 (top panel in A) to 4 *k_B_T* (bottom panel in A). For reference, the *σ* = 0 case is given in each plot (green). With increasing *σ* the oscillations become increasingly dampened, but still nicely reproduced by the nucleosome gas model without landscapes but renormalized parameters *ε*_eff_ and *w*_eff_ (given in each left panel). The panels in (B,D) show the errors *δ*^2^(tot), *δ*^2^(〈*n*(*x*)〉) and δ^2^(〈*n*_2_(*d*)〉) as a function of the landscape amplitude *σ* (see text). The magenta line in the total error plots indicates our cutoff for a good fit, namely *δ*^2^ = 0.003. All considered examples satisfy this criterion.

**FIG. S3:**
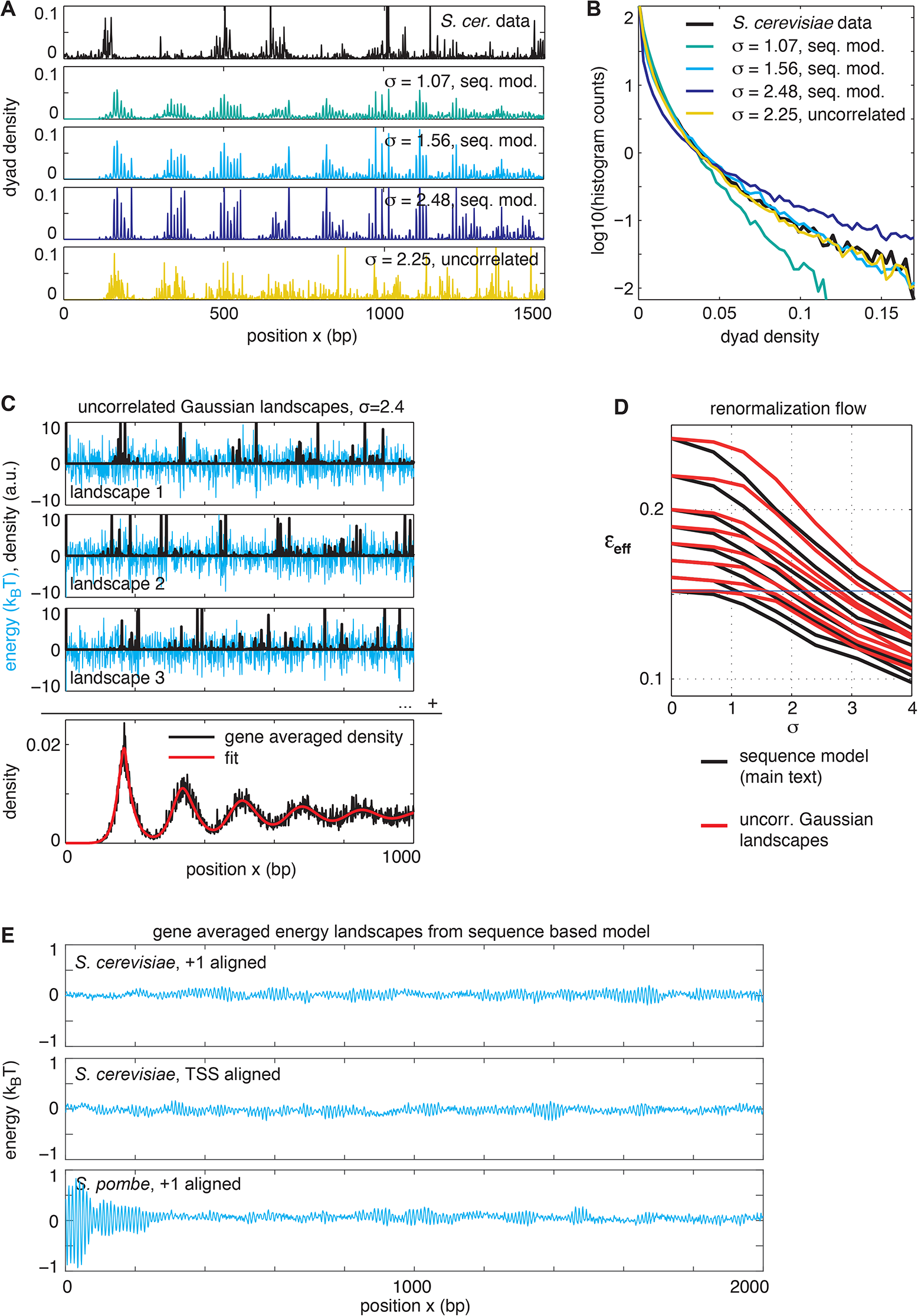
Related to Fig. 3. (A) Examples of nucleosome densities as measured experimentally and predicted by the soft-core nucleosome model on energy landscapes from different models and with various amplitudes *σ*. (B) Histogram of nucleosome density occurrences (see text for details). (C) Uncorrelated Gaussian landscapes where the energy at each basepair is drawn from a Gaussian distribution with a given standard deviation *σ* and corresponding nucleosome densities according to the nucleosome gas model. (D) Parameter flow for the DNA elasticity based energy model used in the main text (black) and uncorrelated Gaussian landscapes (red). (E) Averaged energy landscapes as predicted by the used DNA sequence based model for our sets of *S. cerevisiae* and *S. pombe* genes. The absence of a correlation between the oscillatory gene averaged nucleosome density and the gene averaged landscapes shows that there are no sequence encoded positioning cues on average. This is true in both alignments and both species. Note that this finding does not preclude strong sequence encoded positioning on individual genes.

**FIG. S4:**
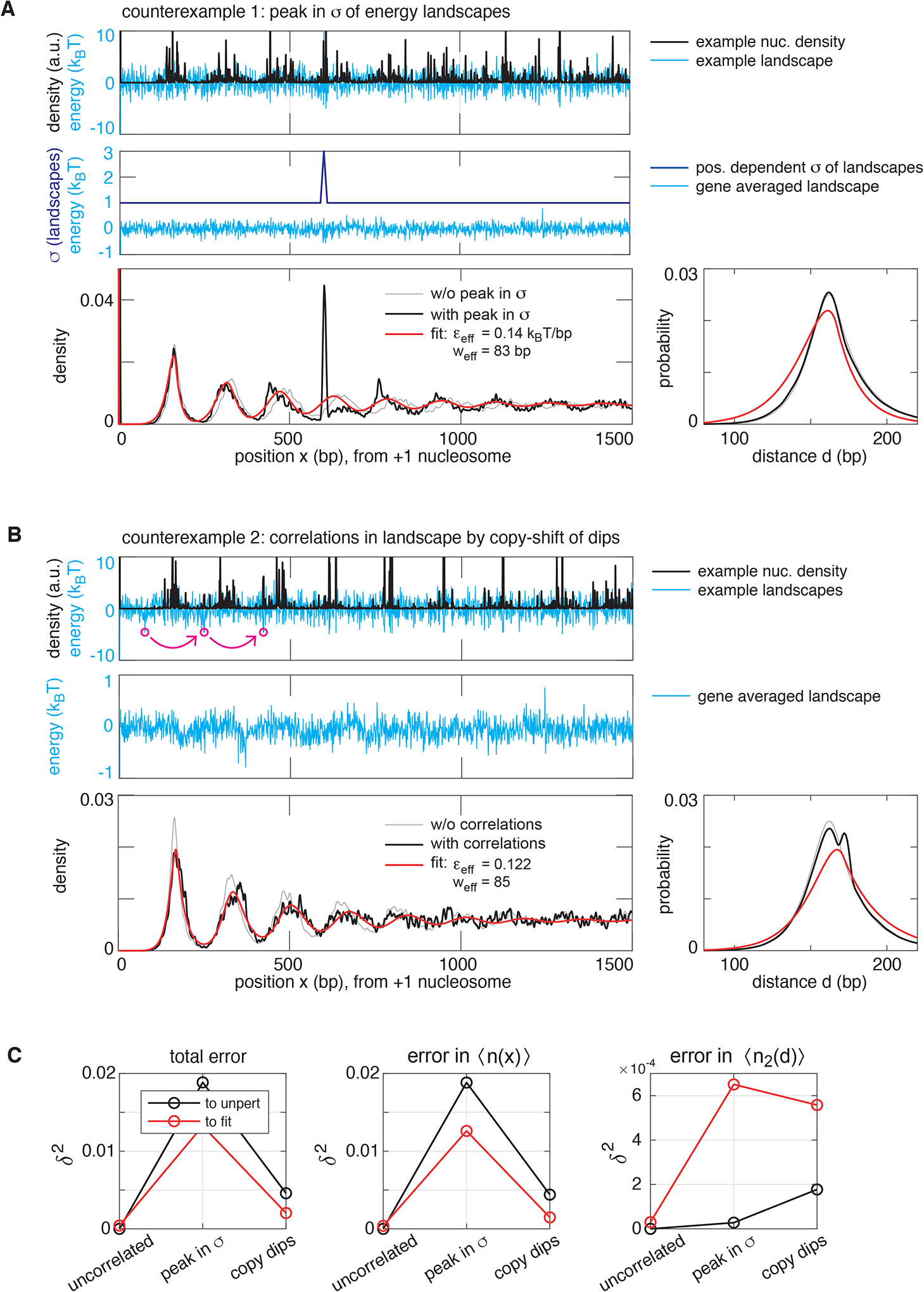
Related to Fig. 3. Examples of positioning landscapes that are incompatible with the nucleosome gas model. (A) Top: examplary random energy landscape (cyan) where the energy at position *x* is drawn from a Gaussian distribution with standard deviation *σ*(*x*). Corresponding nucleosome density in black. Middle: we used *σ*(*x*) = 1 except for a region between bp 590 and 610, where *σ* ramps up to 3 (dark blue). In the landscape averaged over 100 realizations (cyan) the peak in *σ*(*x*) averages out. Bottom: The averaged density has a strong maximum in the peak region (black), which is incompatible with the nucleosome gas (red). (B) Top: We introduce spatial correlations into the landscape by repeating values lower than 2.8 *k_B_T* further downstream, namely shifted by 172, 173, 174 and 345, 346, 347 bp (magenta circles show an example). Middle: In the gene averaged landscape these correlations are barely visible. Bottom: The spacing distribution reflects the correlations in a second peak at the shift of 173 bp, which is incompatible with the nucleosome gas (red). (C) The fit errors show that the discrepancy between data and fit (red) is almost as large as between unperturbed and perturbed data (black). Hence we consider those positioning mechanisms as incompatible with the nucleosome gas model.

**FIG. S5:**
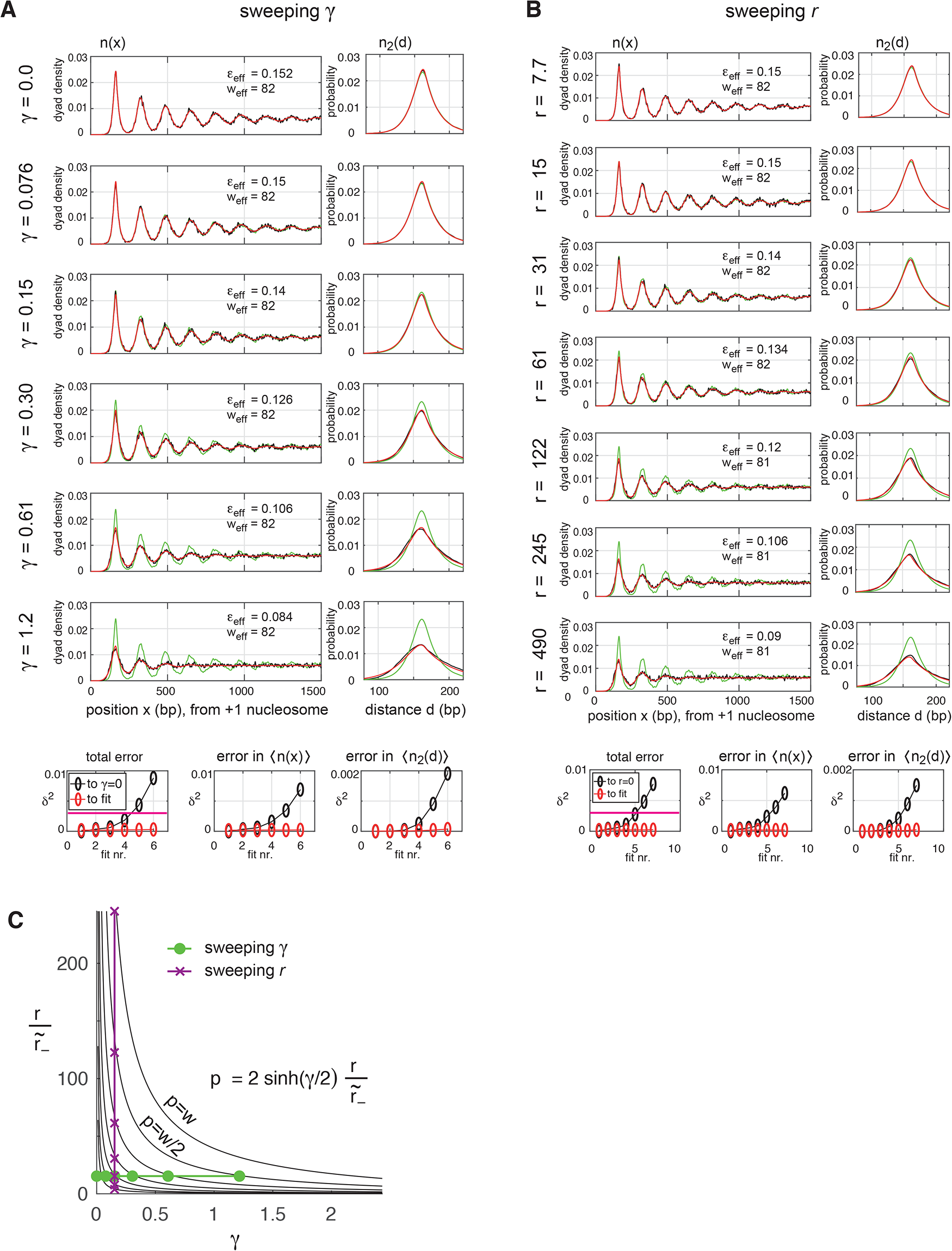
Related to Fig. 4. Fitting the nucleosome gas model to data from remodeler mediated nucleosome sliding. Time averages of the nucleosome density *n*(*x*) and spacing distribution *n*_2_(*d*) in the presence of remodeler mediated directional sliding (black) as described in the main text. The remodeler processivity is varied in two ways, namely by sweeping the bias parameter *γ* (panel A) and the basal activity *A* (panel B). See Table S1 for all parameters. Red: Best fit to data. Green: quantities for *A* = 0 as a reference. The magenta line in the total error plots indicates our cutoff for a “good fit”, namely δ^2^ = 0.003. All considered examples satisfy this criterion. (C) Overview over the used values for *A* and *γ*. The curves are lines of constant remodeler processivity *p*.

**FIG. S6:**
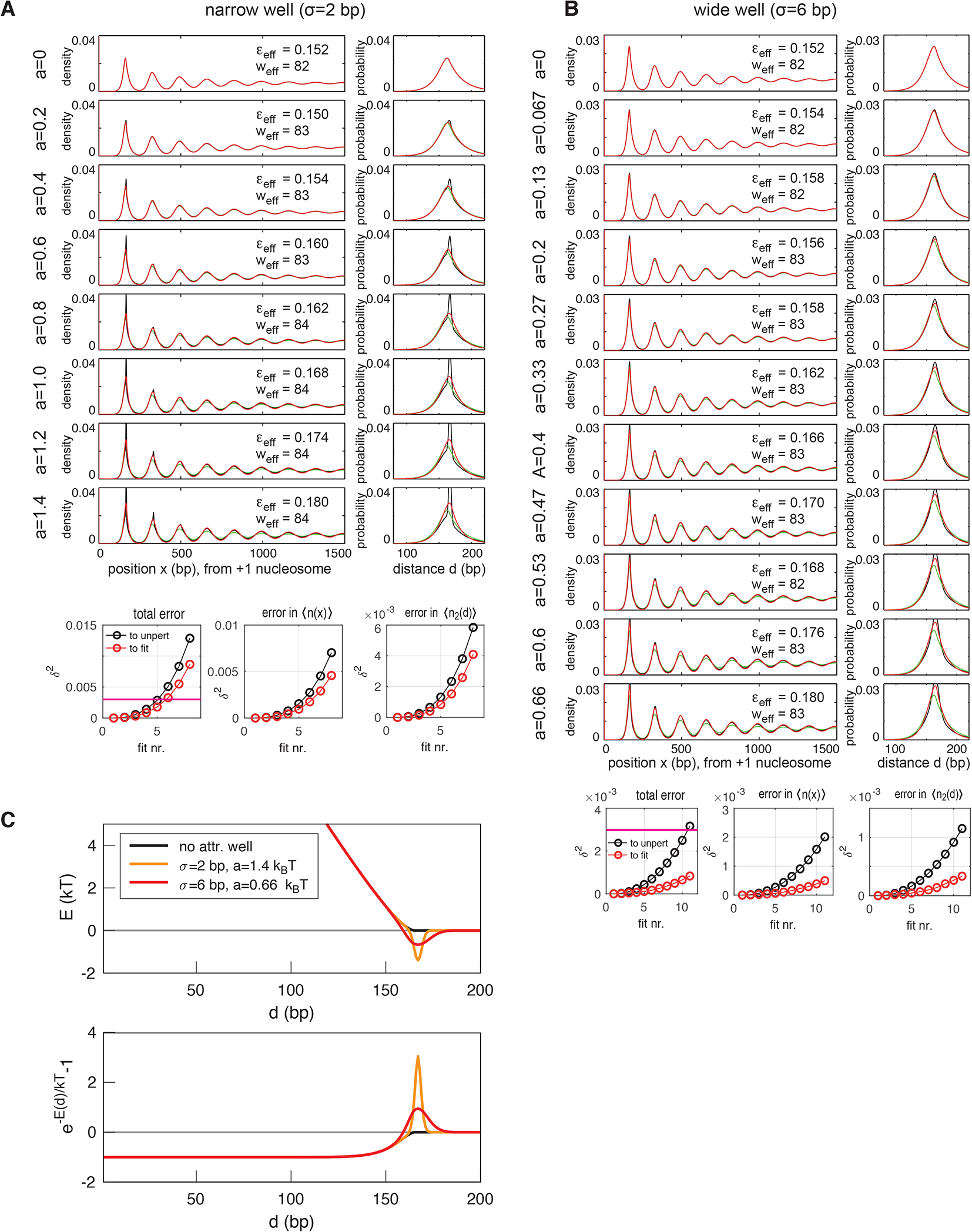
Related to Fig. 5. Fitting the nucleosome gas model to data from remodeler mediated attraction and spacing. (A,B) Simulated nucleosome positioning data (black) for remodeler mediated attraction and spacing for increasing remodeler activity parametrized by the potential well area (see text) for a narrow well with *σ* = 2bp (A) and a wider well with *σ* = 6 bp (B). For the narrow well the fit by the nucleosome gas model (red) is not good, since the discrepancy between the data and the fit is almost as large as between perturbed and unperturbed data (bottom panels). For a wider well fit error is much lower, indicating better compatibility with the nucleosome gas model. The magenta line in the total error plots indicates our cutoff for a “good fit”, namely *δ*^2^ = 0.003. For the narrow well not all considered examples satisfy this criterion, for the wide well all do. (C) Top: example nucleosome interaction potentials for our effective equilibrium description of remodeler mediated attraction and spacing in terms of adding attractive potentials wells to the soft-core repulsive potential. Bottom: corresponding deviations of the Boltzmann factor from unity which appear in the integrand of the second virial coefficient, Eq. (22).

**FIG. S7:**
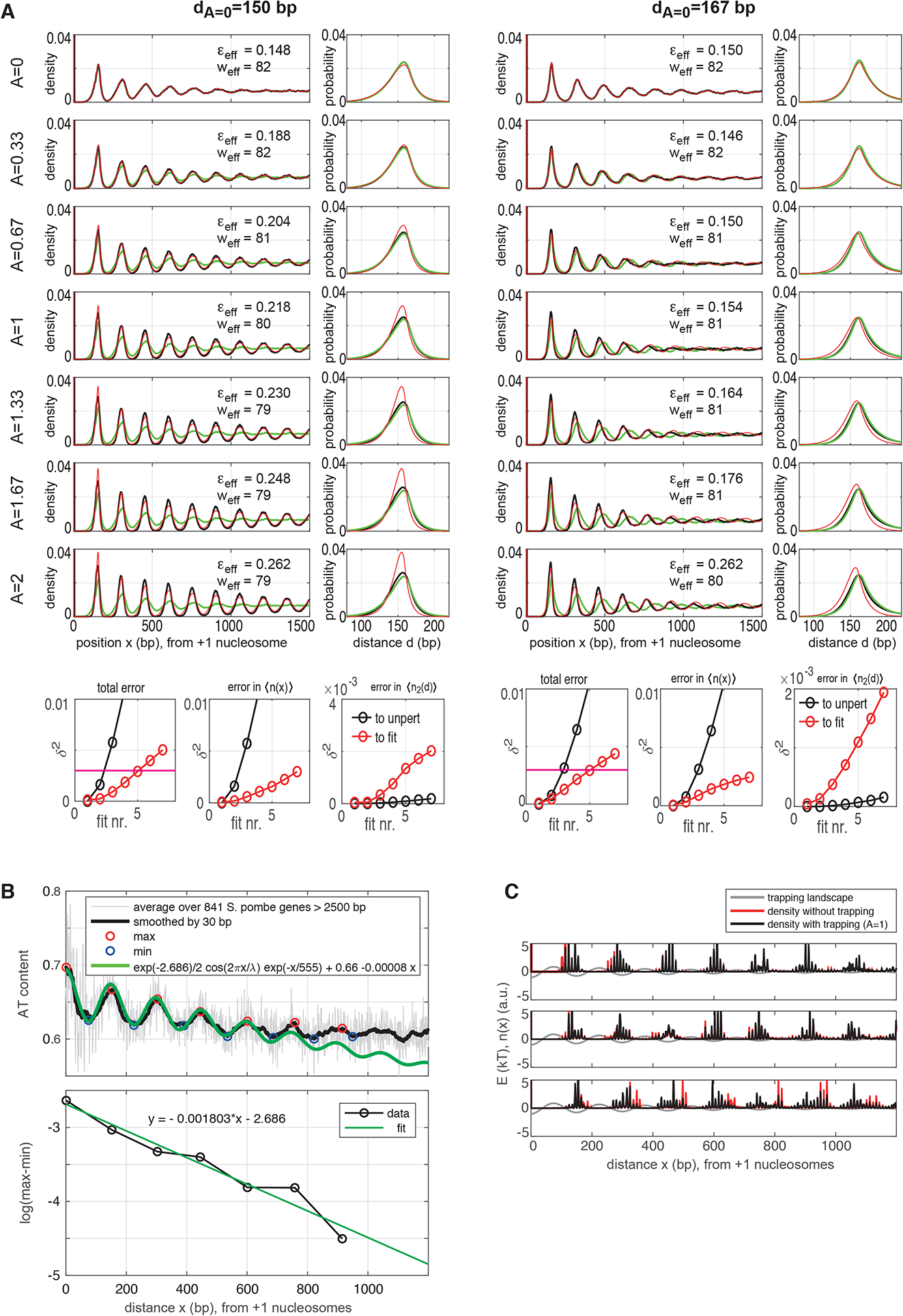
Related to Fig. 7. Fitting the nucleosome gas model to data from a model for trapping on AT rich sequences. (A,B) Simulated nucleosome positioning data (black) on gene-specific energy landscapes superimposed by a decaying oscillatory landscape to account for trapping on AT rich sequences on average (see text for details). From top to bottom the maximal amplitude *A* of the trapping landscape is increased. Red: Best fit to data. Green: quantities for *A* = 0 as a reference. Left: the period of the trapping landscape equals the density peak-to-peak distance (both are 150 bp, like in *S. pombe*). Right: the period of the trapping landscape is 150 bp (like *S. pombe*), but the density peak-to-peak distance without trapping is 167 bp (like *S. cerevisiae*). Bottom panes: fit errors. The magenta line in the total error plots indicates our cutoff for a “good fit”, namely *δ*^2^ = 0.003. For too strong trapping the fits fail to meet this criterion. (B) AT content in *S. pombe* genes. The max-to-min difference decays exponentially with increasing distance from the +1 nucleosome. (C) Density on example genes computed with and without a trapping landscape in addition to gene specific energy landscapes derived from a DNA elasticity based model (see main text). The trapping landscape is our model for positioning of nucleosomes on AT rich sequences that is observed in *S. pombe* data. On individual genes the influence appears rather small, but when averaged over many genes the resulting oscillations are markedly more pronounced, which leads to an effective stiffening (see main text).

